# NatB modulates Rb mutant cell death and tumor growth by regulating EGFR/MAPK signaling through the N-end rule pathways

**DOI:** 10.1101/590349

**Authors:** Zhentao Sheng, Wei Du

## Abstract

Despite the prevalence of N-terminal acetylation (Nt-acetylation), little is known of its biological functions. In this study, we show that NatB regulates Rb mutant cell survival, EGFR/MAPK signaling activity, and EGFR signaling-dependent tumor growth. We identify Grb2/Drk, MAPK, and PP2AC as the key NatB targets of EGFR pathway. Surprisingly, NatB activity increases the levels of positive pathway components Grb2/Drk and MAPK while decreases the levels of negative pathway component PP2AC despite these proteins have the same first two amino acids that are recognized by NatB and N-end rule pathways. Mechanistically, we show that NatB regulates Grb2/Drk protein stability through its N-terminal sequences and that Grb2/Drk and MAPK are selectively degraded by the Arg/N-end rule E3 ubiquitin ligase Ubr4, which targets proteins with free N-terminus. In contrast, PP2AC is selectively degraded by the Ac/N-end rule pathway E3 ubiquitin ligase Cnot4 that targets proteins with acetylated N-terminus. These results reveal a novel mechanism by which NatB-mediated Nt-acetylation and N-end rule pathways modulate EGFR/MAPK signaling by inversely regulating the levels of positive and negative components. Since mutation or overexpression that deregulate the EGFR/Ras signaling pathway are common in human cancers and NatB subunits are significant unfavorable prognostic markers, this study can potentially lead to the development of novel therapeutic approaches.

**Significance Statement:** Nt-acetylation is often regarded as a constitutive, irreversible, and static modification that is not suited to serve regulatory functions. Our observation that Nt-acetylation by NatB coordinately regulate the levels of positive and negative components of the EGFR/MAPK pathway show that Nt-acetylation and N-end rule pathways can play important roles regulating important signaling pathways. As Acetyl-CoA level, which is influenced by cell metabolism, can be rate limiting for Nt-acetylation, our results also suggest a potentially new mechanism by which cellular metabolic status can regulate growth factor signaling.

## Introduction

N-terminal acetylation (Nt-acetylation), which is observed in majority of the proteins in eukaryotes, involves the transfer of an acetyl group from acetyl-CoA to the α-amino group of a protein. Nt-acetylation is catalyzed by the N-terminal acetyltransferases (NATs). There are six NATs, NatA-F, in higher eukaryotes with different substrate specificity [1]. Nt-acetylation neutralizes the positive charge and alters chemical properties of the N terminus of the protein. At biochemical level, Nt-acetylation has been shown to affect localization, interaction, and/or degradation of specific proteins. However, despite the widespread nature of this modification [2], the biological function of NAT and their target Nt-acetylation is still largely unknown.

Studies of engineered β-galactosidases in yeast has established that the Nt-residuals play critical roles in determining protein stability through the N-end rule pathway [3, 4]. Two branches of the N-end rule pathway have been identified: the Arg/N-end rule pathway and the Ac/N-end rule pathway [3, 5]. In Arg/N-end rule pathway, nonacetylated destabilizing Nt-residues were recognized by the N-recognins, which induce target protein ubiquitination and degradation. N-recognins for the Arg/N-end rule pathway were found to be the UBR-box containing E3 ubiquitin ligases conserved from yeast to mammals. The UBR box is essential for the binding of the N-terminal destabilizing residues and the recognition of the positively charged N-terminal NH3+ group is critical [6]. Interestingly, of the seven UBR box containing E3 ubiquitin ligases found in mammalian genomes, only four (Ubr1, Ubr2, Ubr4, and Ubr5) were found to be N-recognins [3]. On the other hand, cellular proteins with Nt-acetylated residues will be recognized by the Ac/N-recognins, which will target the ubiquitination and degradation of Nt-acetylated proteins through the Ac/N-end rule pathway. Two Ac/N-recognins Doa10 and Not4 have been identified in yeast. The Doa10 homolog Teb4 has been shown to function as a mammalian Ac/N-recognins [7], suggesting that the Ac/N-end rule pathway is also conserved. The recognition by Ac/N-recognins was proposed to be conditional, influenced by protein folding and protein complex formation. The conditional nature of degradation by the Ac/N-end rule pathway provides a mechanism for protein quality control and balance the levels of subunits in a protein complex [5].

Because Nt-acetylation is believed to be largely co-translational and without a corresponding deacetylase, Nt-acetylation is often regarded as a constitutive, irreversible, and static modification that is not suited to serve regulatory functions. In addition, as studies of N-end rule pathways have mostly been carried out in yeast using a reporter protein, very little is known about the regulation of endogenous proteins from multicellular organisms by the two branches of N-end rule pathways. Indeed, it is not known whether Nt-acetylation/N-end rule pathways can regulate the signaling output of major signaling pathways. As there are many potential Nt-acetylation targets in a major signaling pathway, it is not clear whether Nt-acetylation/N-end rule pathways will regulate different targets that serve positive or negative functions coordinately, which would be expected if Nt-acetylation/N-end rule pathways were to play an important role in regulating a signaling pathway.

*Drosophila* provides a genetically tractable model with well-conserved genes and important signaling pathways with humans. In *Drosophila* developing eye discs, inactivation of the fly Retinoblastoma (Rb) homolog Rbf induces high levels of cell death specifically near the morphogenetic furrow (MF) [8], a region in eye disc where anterior asynchronous progenitor cells arrest in G1 and initiate photoreceptor differentiation. This cell death is mediated by Hid induction and are blocked by mutations that can increase EGFR signaling [9-11]. EGFR signaling is activated in the MF and posterior region of the developing eye discs and plays critical roles in preventing cell death in addition to its role in regulating cell cycle and differentiation [12]. Interestingly, reducing EGFR signaling by mutations that function upstream of MAPK preferentially increased apoptosis of *rbf* mutant cells in posterior eye discs [11, 13]. In contrast, mutation of *rno*, which functions downstream of MAPK and regulates the nuclear output of EGFR signaling [14], did not affect apoptosis of *rbf* mutant cells. This is consistent with the reported direct regulation of Hid by MAPK [15]. Taken together, these results suggest that *rbf* mutant cells are more sensitive to reduced level of EGFR/MAPK signaling and that the *rbf* synthetic lethal screens [13, 16, 17] can identify additional modulators of EGFR signaling. In this study, we report new alleles of NatB subunits in *rbf* synthetic lethal screen and identify a novel mechanism by which NatB modulates EGFR signaling by regulating the levels of positive and negative components of the pathway coordinately through the two branches of the N-end rule pathway.

## Results

### Inactivation of NatB subunits induced synergistic cell death with loss of *rbf* in developing *Drosophila* tissues

In a genetic screen to identify Rbf synthetic lethal mutations on *Drosophila* chromosome 3R, we identified four mutants (*AB*, *AE*, *CJ*, and *EQ*) that fall into the same complementation group. While clones of FRT control or single mutant (paled patches) were readily detected in adult eyes, double mutant clones of *rbf* and these mutants were not detectable (Fig. 1A-D, Fig. S1I-J). To better visualize the loss of double mutant tissues, we generated single or double mutant clones in the *Minute* background, which allows large mutant clones to be generated due to competitive growth advantage [18]. Indeed, large white patches of *AE*, *EQ*, *AB*, and *CJ* single mutant clones were observed in *Minute* background (Fig. S1A, C, E, G). Interestingly, most of the white patches were also lost in conjunction with *rbf* mutation, leading to the development of much smaller adult eyes (Fig. S1B, D, F, H). These results strongly support the notion that the identified mutants induce synthetic lethality in conjunction with *rbf* mutation.

**Fig. 1.**
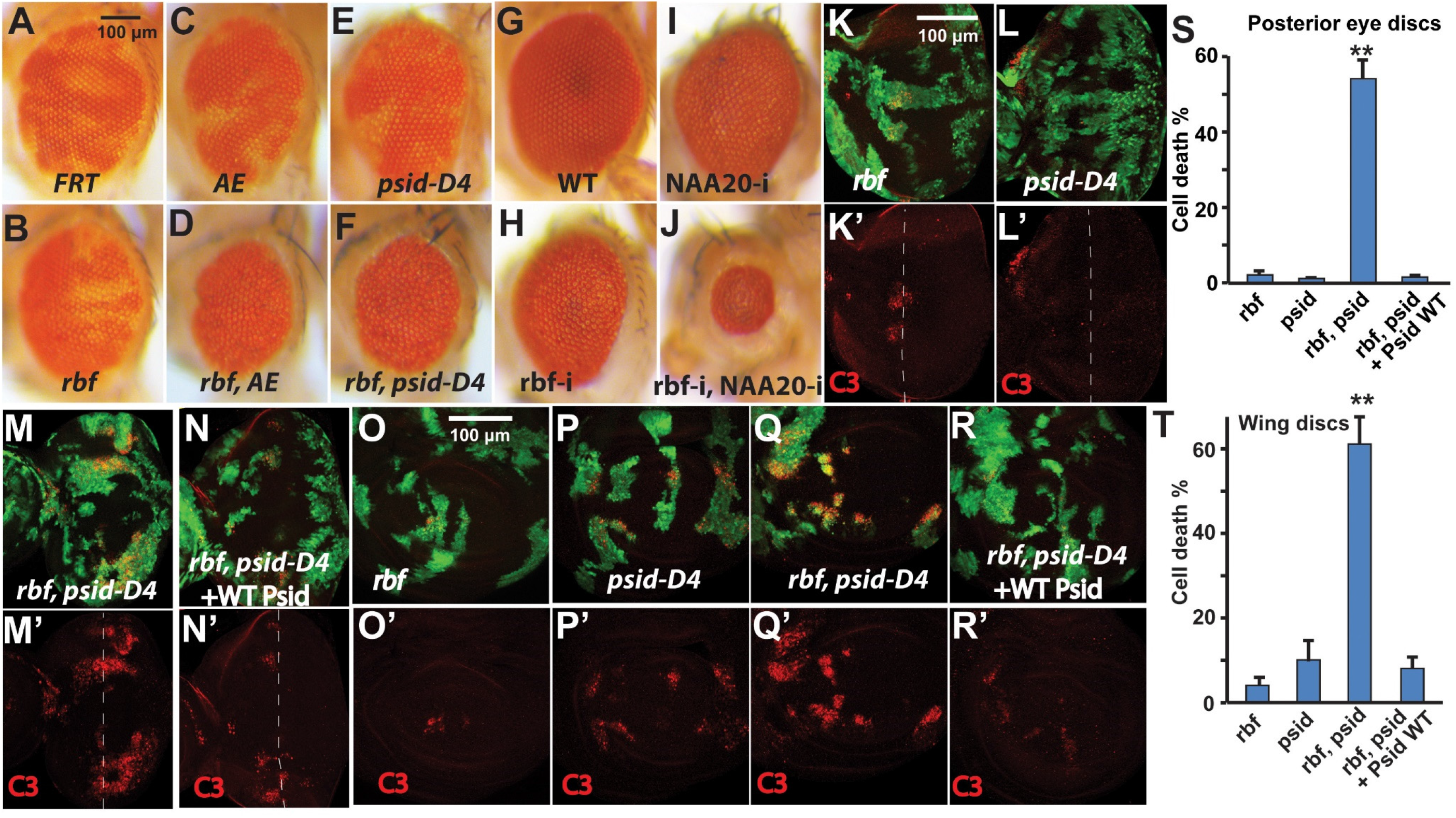
Inactivation of NatB subunits, Psid or NAA20, induced synergistic cell death with loss of *rbf* in *Drosophila*. (A-F) Adult eyes with clones of WT control (A), *rbf*, *psidin* single or double mutation (B-F). *AE* and *psid-D4* (*psid-55D4*) are two different *psidin* alleles. Mutant clones are marked by lack of red pigment. (G-J) Adult eyes with single or double RNAi of Rbf or NAA20 were shown. (K-T) Levels of cell death in 3^rd^ instar eye discs (K-N’) or wing discs (O-R’) with *rbf*, or *psidin*, or *rbf psidin* double mutant clones are shown. Mutant clones are marked by GFP expression and cell death are determined by caspase 3 staining shown in red. Dash line indicates the position of furrow region in developing eye discs (K-N). Levels of cell death in mutant clones located in posterior eye discs and wing discs were quantified and the average and standard deviation were shown in (S) and (T), respectively (N≥6 for each genotype). In both posterior eye discs (K-M, S) and wing discs (O-Q, T), the levels of cell death were significantly increased in *rbf, psidin* double mutant clones in comparison to the single mutant clones. Expression of WT Psid rescued the cell death in *rbf, psidin* double mutant clones in both eye (M-N, S) and wing discs (Q-R, T). ** indicates statistically significant differences (p<0.0001, t test) between double and single mutant clones or between double mutant clones with or without WT Psid rescue. The black or white bars indicate 100 µM. In this and all the subsequent figures, discs are orientated posterior to the right, genotypes of mutant clones and anti-staining are indicated on the figure.

Genetic mapping using 3R deficiencies was carried out. We found that the *AE* complementation group mutants map to the 92B4-C1 cytological region. The following results demonstrate that the *AE* complementary group mutants are alleles of *psid.* First, both of the two existing *psid* alleles, *psid-D4* (*psid^55D4^*) and *psid-D1* (*psid^85D1^*) [19, 20], failed to complement the *AE* group mutants; Second, both *psid-D4* and *psid-D1* showed loss of mutant clones in the presence of *rbf* mutation, similar to that of *AE* and *EQ* mutants (Fig. 1C-F, Fig. S1I-L).

To demonstrate that *psid* mutation induced synthetic lethality with *rbf*, we used anti activated caspase 3 antibody to determine cell death in the developing discs. *rbf* mutant cells showed significantly increased cell death in the morphogenetic furrow (MF) region but not much in the posterior of the developing eye disc [8, 9] (Fig. 1K-K’). While *psid* single mutant clones did not increase cell death in posterior eye discs (Fig. 1L-L’), significant increased cell death was observed in *rbf, psid* double mutant clones in posterior eye discs (Fig. 1M-M’, 1S). Importantly, expression of WT Psid rescued the observed synergistic cell death in posterior eye disc (Fig. 1N-N’, 1S), demonstrating that the increased cell death depended on the loss of Psid function. Furthermore, synergistic cell death of the *rbf, psid* double mutant cells, which can be rescued by expression of WT Psid, was also observed in the wing discs (Fig. 1O-R’, 1T), indicating that the cell death effects of *rbf psid* double mutants are not specific to the eye discs.

Psid is the regulatory subunit of N-terminal acetyltransferase B (NatB). It binds to the catalytic subunit NAA20 to catalyze the addition of acetyl group to the N terminus of proteins that start with MD, ME, MN, or MQ. Knockdown of NAA20 by RNAi in conjunction with *Rbf* RNAi also led to significantly decreased adult eye sizes (Fig. 1G-J), which was correlated with significantly increased cell death in eye/antenna discs (Fig. S1M-O). Taken together, these data suggest that inactivation of NatB induced synergistic cell death with loss of *rbf*.

### NatB regulates MAPK activation at multiple points downstream of EGFR

Our previous studies suggest that mutations such as *TSC2* and *axin* induce synergistic cell death with *rbf* most strongly in anterior eye discs due to induction of excessive cellular stress [16, 17, 21]. In contrast, mutations that cause deficiency in EGFR/MAPK signaling induce synergistic cell death with *rbf* mainly in posterior eye discs [13]. The observed *psid, rbf* synergistic cell death in posterior eye discs prompted us to determine whether *psid* mutation affect EGFR/MAPK signaling. EGFR signaling is upregulated in posterior eye discs, which play important roles regulating cell cycle, cell survival, and differentiation [12, 22]. Indeed, *psid* mutation significantly decreased EGFR signaling as shown by reduced Aos-lacZ reporter expression and reduced level of active Diphosphorylated ERK (pERK) in *psid* mutant clones (Fig. 2A-B”, Fig. S2A-C’). Furthermore, expression of WT Psid restored the pERK levels (Fig. 2C-C”). Interestingly, expression of the phosphomimetic Psid mutant (Psid^S678D^), which mutated a conserved Ser that was shown to be phosphorylated in human MDM20 [23], failed to rescue (Fig. 2D-D”), while expression of the nonphosphorylatable Psidin^S687A^ mutant did rescue (Fig. 2E-E”). As the Psid^S678D^ mutant is defective in binding to NAA20 while the Psid^S687A^ mutant retains the ability to bind NAA20 [20], our results suggest that the reduced MAPK activation in *psid* mutant clones is mediated by reduced NatB activity. In support of this, knockdown of NAA20 in eye discs also significantly reduced EGFR signaling as shown by reduced Aos-lacZ reporter expression (Fig. 2F-F’) and reduced pERK levels (Fig. S2D-D’).

**Fig. 2.**
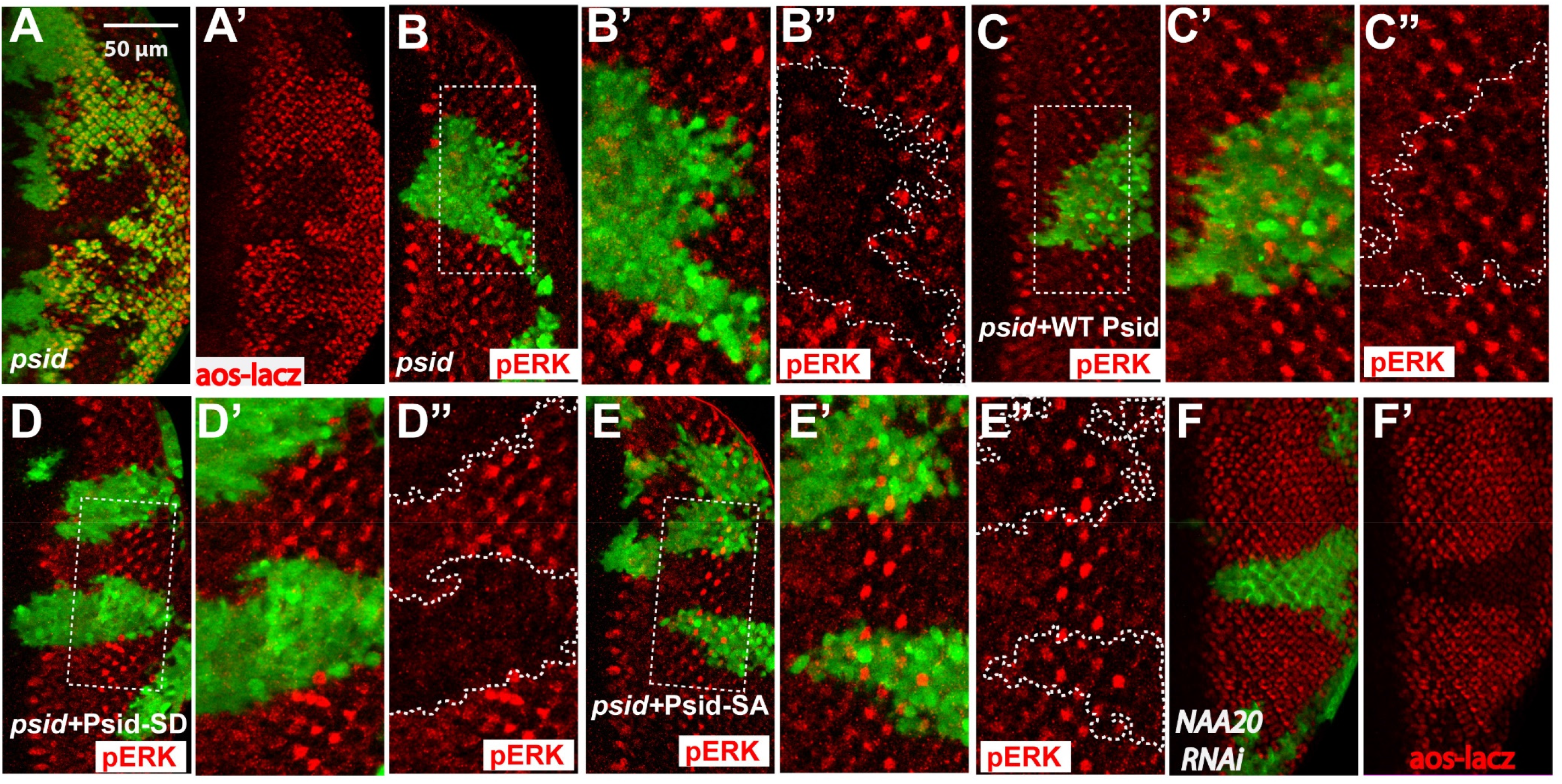
NatB was required for the elevated EGFR signaling in posterior eye discs. (A and A’) *psid-D4* clones in posterior of eye discs were marked by the absence of GFP. The level of EGFR signaling, as shown by the expression of aos-lacZ reporter detected by anti β-galactosidase staining (red), was significantly decreased in *psid-D4* clones. (B-E”) MARCM clones of *psid-D4* in the posterior of eye discs were marked by GFP expression. EGFR signaling, as shown by the level of pERK, was significantly decreased in *psid-D4* clone (B-B”), and was restored by expressing WT Psid (C-C”) or the nonphosphorylatable Psid^SA^ (S678A) (E-E”) but not the phosphomimetic Psid^SD^ (S678D) mutant (D-D”). (F and F’) NAA20 RNAi clones marked by GFP expression also decreased EGFR signaling as shown by decreased expression level of aos-lacZ.

Ligands for EGFR signaling was released by the developing photoreceptor cells, particularly the R8 photoreceptor cells in the developing eye discs. Knockdown NAA20 slightly delayed the formation R8 equivalence groups and the differentiation of R8 and additional photoreceptor cells (Fig. S2E-F’). However, the differentiation of photoreceptor cells was not blocked in the posterior where decreased EGFR signaling was observed. These observations suggest that NatB affects EGFR signaling not simply through affecting photoreceptor differentiation. To further investigate how *psid* mutation inhibits EGFR signaling, we determined the effect of *psid* mutation on MAPK activation induced by expressing activated EGFR, Ras, or Raf in *psid* mutant or WT control cells using MARCM approach. Interestingly, activated EGFR significantly increased pERK levels in WT control cells but not in *psid* mutants (Fig. 3A-B’). In contrast, activated Raf-induced pERK levels was not obviously inhibited by *psid* mutation (Fig. 3E-F’, Fig. S3C-D’). On the other hand, even though activated Ras-induced pERK was largely inhibited by *psid* mutation, activated Ras can induce slightly elevated pERK levels in *psid* mutants (Fig. 3C-D’, Fig. S3A-B’). These results suggest that *psid* mutation blocks MAPK activation at multiple points downstream of activated EGFR and upstream of activated Raf (Fig. 3L). To demonstrate that the observed inhibition of activated EGFR-induced pERK level is mediated by the loss of NatB activity, the ability of expressing WT or mutant Psid proteins to rescue EGFR-induced pERK was determined. The WT and S687A mutant form of Psid that can bind NAA20 were able to restore EGFR-induced pERK levels in *psid* mutant clones (Fig. 3G-H’) while the NAA20 binding defective Psid^S678D^ mutant failed to rescue (Fig. 3I-I’). Furthermore, knockdown of NAA20 by RNAi also strongly inhibited activated EGFR-induced pERK (Fig. 3J-K’). Taken together, these results suggest that NatB modulates the activities of multiple targets that function between EGFR and Raf to regulate MAPK activation.

**Fig. 3.**
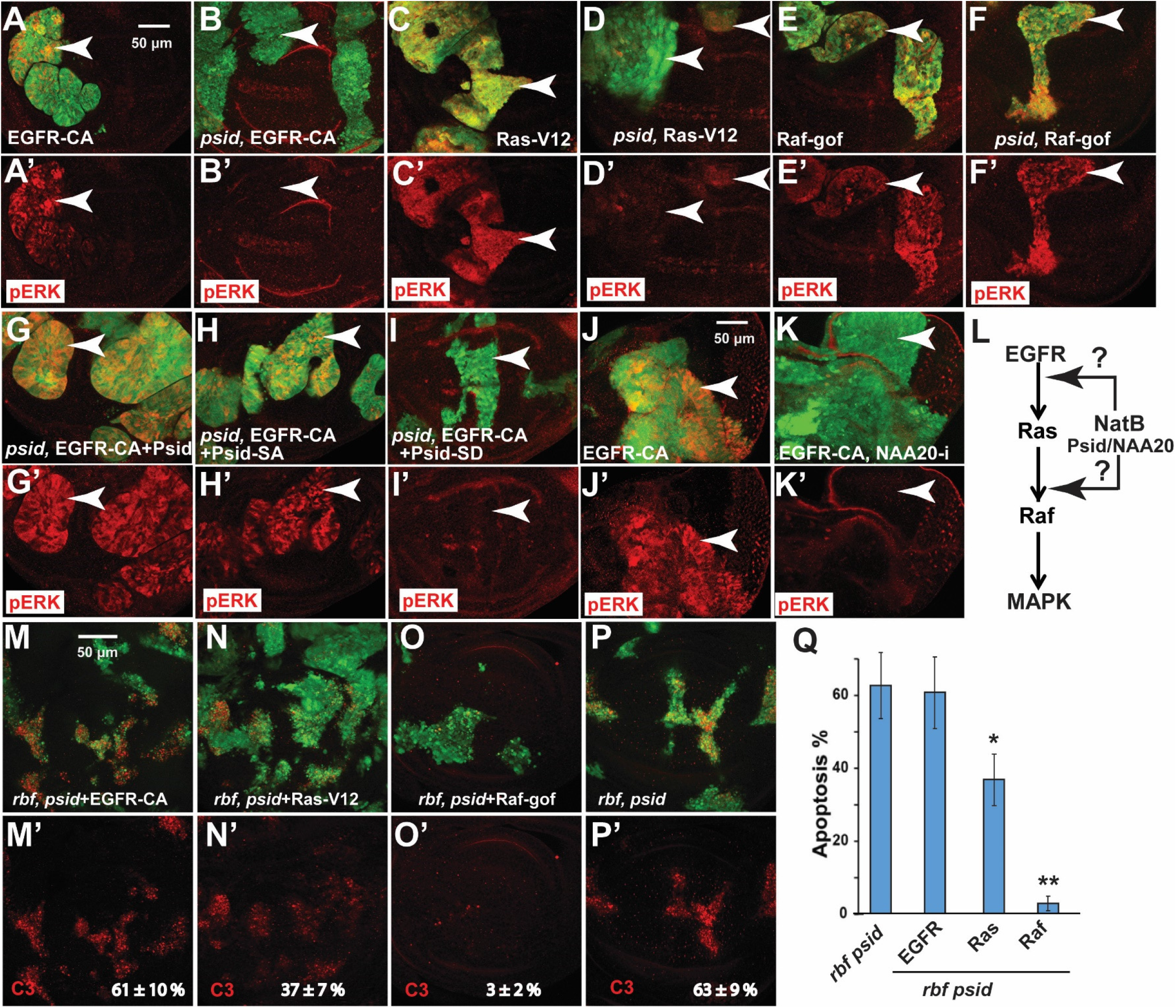
Inactivation of Nat B subunits inhibited MAPK activation at multiple points downstream of EGFR. (A-F’) The effects of *psidin* (*psid-D4*) mutation on the MAPK activation (pERK staining, red) induced by expressing activated EGFR, Ras, or Raf in the wing discs are shown. *psid-D4* mutant clones with UAS transgenes were marked by GFP expression. Activated EGFR-induced pERK was completely inhibited by *psid-D4* mutation (A-B’). In contrast, activated Ras-induced pERK was partially inhibited by *psid-D4* mutation (C-D’) while activated Raf-induced pERK was not significantly inhibited (E-F’). (G-I’) Inhibition of activated EGFR-induced pERK in *psid* (*psid-D4*) mutant clones was rescued by expressing WT Psid (G) or the nonphosphorylatable Psid^SA^ (S678A) (H) but not the phosphomimetic mutant Psid^SD^ (S678D) (I). (J-K’) Using EyeFlp-CoinGal4, activated EGFR-induced pERK (J) was largely inhibited by NAA20 RNAi in eye disc (K). (L) A diagram that summarizes the main points that NatB regulated EGFR/MAPK signaling. (M-Q) Cell death in *rbf, psidin* double mutant clones with different UAS-transgene expression in wing discs were determined by C3 staining (red). The double mutant clones were marked by GFP expression. While activated EGFR (M) did not inhibit cell death observed in *rbf, psid* double mutant clones (P), activated Ras partially inhibited cell death (N), and activated Raf inhibited cell death to background levels (O). (Q) The level of cell death in mutant clones located in wing discs were quantified and the average and standard deviation were shown. The statistically significant difference between triple and double mutant clones were showed (N≥6 for each genotype; ** indicates P<0.0001, * indicates P<0.005, t test).

To determine whether inhibition of EGFR signaling induced MAPK activation contributed to the synergistic cell death of *rbf, psid* double mutant cells, we tested the ability of activated EGFR, Ras, or Raf to rescue *rbf, psid* double mutant cell death. While activated EGFR failed to rescue *rbf, psid* cell death, activated Ras induced partial rescue and activated Raf rescued cell death to background levels (Fig. 3M-Q). These results support the notion that reduced MAPK activity in *psid* mutant clones contributed to the synergistic cell death with *rbf*.

### *psid* mutation inhibited activated EGFR-induced tumor growth

Activation of EGFR signaling in conjunction with *scrib* mutation induces cephalic and male gonadal tumors when Ey-FLP was used to induce MARCM clones [24-26]. Interestingly, *psid* mutation significantly inhibited activated EGFR-induced tumor growth in both regions (Fig. S3E, Yellow and white arrowheads, tumor cells were marked by GFP). The gonadal tumor is initiated by Ey-FLP induced *scrib* MARCM clone in the ‘terminal body’ (TB) cells at the posterior pole, which grows and spreads to the anterior [26]. Indeed, while the *EGFR^CA^ scrib* mutant cells were highly proliferative as shown by the large numbers of BrdU incorporating cells (Fig. S3G-G’), inactivation of *psid* significantly inhibited the proliferation of *EGFR^CA^ scrib* mutant cells (Fig. S3H-H’). Furthermore, expressing Psid^WT^ or Psid^SA^ but not the NAA20 binding defective Psid^SD^ mutant restored *EGFR^CA^ scrib* tumor growth in *psid* mutant background (Fig. S3F). These results suggest Psid-regulated NatB activity is require for activated EGFR-induce tumor growth.

### NatB stabilizes Drk, which promotes EGFR induced MAPK activation

Since NatB catalyze the addition of acetyl group to the N terminal proteins that starts with MD, ME, MN, or MQ, we analyzed components of the EGFR/MAPK pathway and identified three potential NatB targets, Drk (fly Grb2 homolog), PP2AC (catalytic subunit of PP2A), and Sprouty (Spry), which function between EGFR and Raf. Two of these three proteins, Drk and PP2AC, showed high sequence conservation in the first two amino acid, the N-terminal region, as well as the overall protein. The high level of sequence conservation suggest that the regulation of these two proteins by NatB/N-end rule pathways and the overall function of these proteins are likely conserved. Therefore, we initially focused on Drk and PP2AC.

We first characterized an antibody against Drk [27]. We found Drk RNAi clones decreased Drk protein levels (Fig. 4H-H’), indicating that the anti Drk antibody specifically recognized endogenous Drk protein. Significantly decreased Drk levels were observed in *psid* mutant clones in both eye and wing discs (Fig. 4A-B’, white arrows). Similarly, *psid* MARCM clones also significantly reduced Drk levels (Fig. 4C-C’). In contrast, *psid* mutation did not significantly affect β-gal level expressed from a Drk enhancer trap line (Fig. S4A), suggesting *psid* mutation did not affect Drk expression. Furthermore, expression of Psid^WT^ or Psid^SA^ mutant can rescue Drk levels in *psid* mutant clones while the NAA20 binding defective Psid^SD^ mutant failed to rescue (Fig. 4C-F’). In addition, knockdown of NAA20 using RNAi also significantly decreased Drk levels (Fig. 4G-G’). These results show that inactivation of NatB decreased levels of Drk posttranscriptionally.

**Fig. 4.**
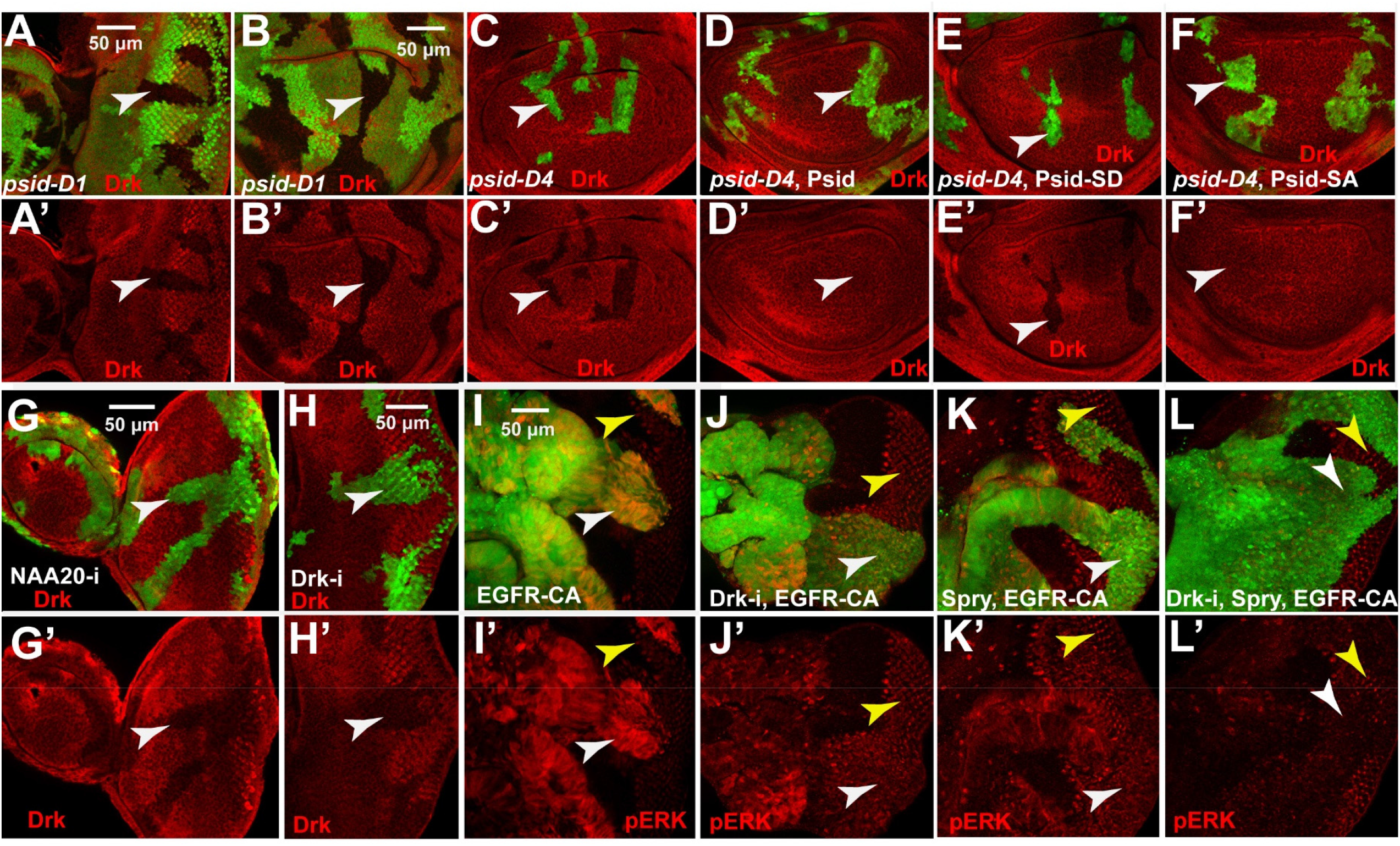
Inactivation of NatB destabilized Drk, which mediated activated EGFR-induced MAPK activation. (A-B) *psid-D1* mutant clones, marked by lack of GFP expression, show decreased Drk level (anti-Drk staining, red) in both eye discs (A) and wing discs (B). (C-F) MARCM clones of *psid-D4* in wing discs, marked by GFP expression, showed decreased Drk level (C) which was rescued by expressing WT Psid (D) or nonphosphorylatable Psid^SA^ (S678A) (F) but not the phosphomimetic mutant Psid^SD^ (S678D) (E). (G-H) GFP positive clones of NAA20 RNAi (G) or Drk RNAi (H) in eye discs, generated with eyeflp-CoinGal4, showed decreased Drk level. (I-L) Drk RNAi (J) reduced the level of activated EGFR-induced pERK (I), which was further inhibited by the expression of Spry (K-L).

To determine the effect of decreasing Drk levels on EGFR signaling, the effect of Drk knockdown on activated EGFR-induced MAPK activation were determined. While activated EGFR induced much higher pERK levels than the endogenous pERK observed in posterior eye discs (Fig. 4I-I’, compare white and yellow arrowheads), Drk-RNAi significantly reduced this EGFR induced pERK levels to that similar to endogenous pERK levels (Fig. 4J-J’, compare white and yellow arrowheads). In addition, expression of Spry in conjunction with knockdown Drk further inhibited EGFR-induced MAPK activation (Fig. 4I-L’). Therefore, reduction of Drk levels can significantly inhibit EGFR-induced MAPK activation.

### Psid regulates Drk through its N-terminal sequences

Since NatB proteins modify Protein N-terminus, which may affect protein stability [3], we directly tested this idea by expressing a GFP protein tagged with Drk N-terminal 10 amino acids (N-Drk-GFP) together with the β-gal control using the MARCM system. The ratio of GFP to β-gal levels were determined in WT or *psid* mutant background and compared. In comparison to the WT control, mutation of either *psid^D1^ or psid^D4^* significantly decreased levels of N-Drk-GFP (Fig. 5A-B”, 5D) but not the WT control GFP that do not have the N-Drk tag (Fig. 5D).

**Fig. 5.**
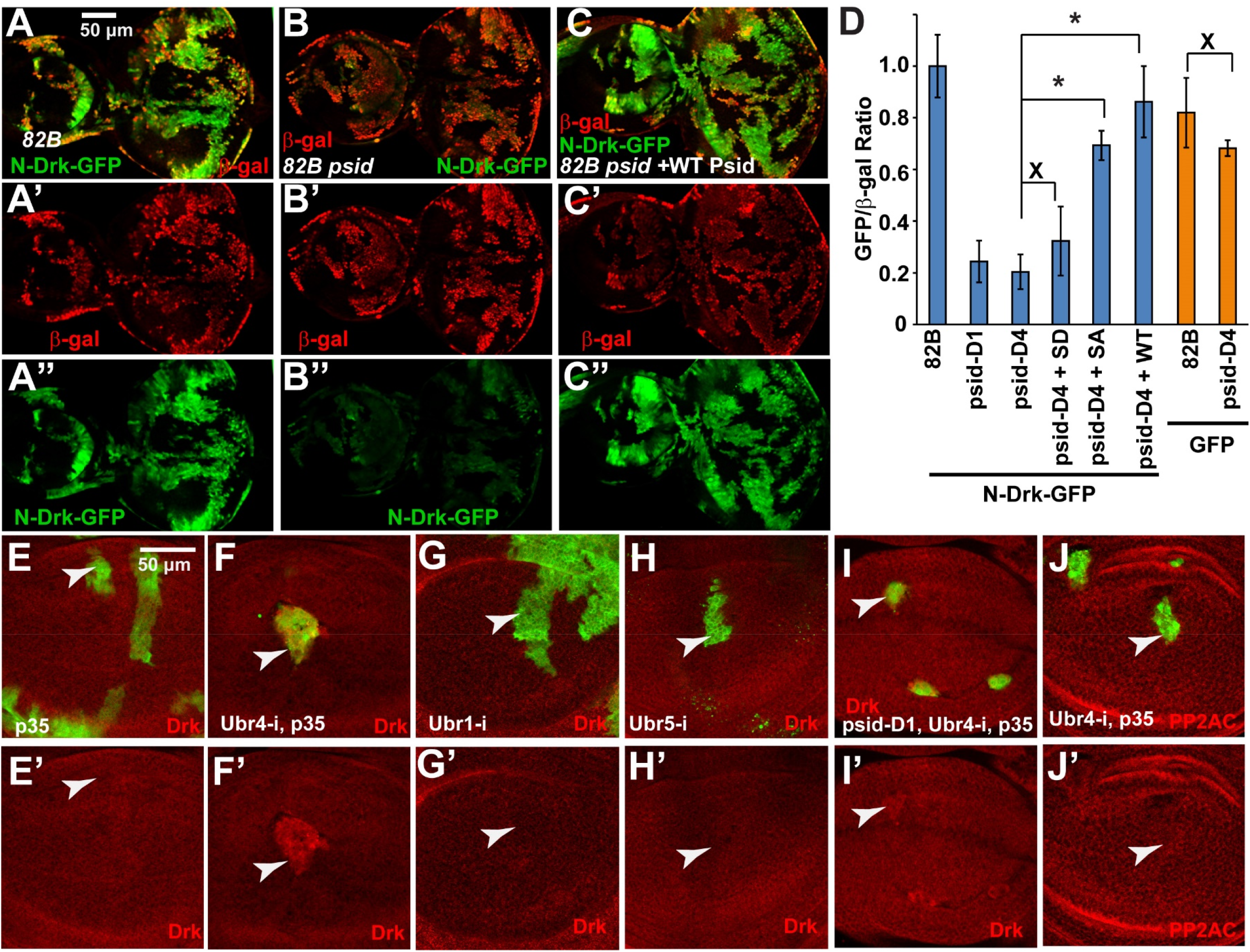
NatB-inactivation decreased Drk levels through its N-terminal sequences, which was regulated by the N-end rule E3 ubiquitin ligase POE/UBR4. (A-C) N-Drk-GFP (GFP tagged with a 10 amino acids N-terminal from Drk, green) and LacZ (β-galactosidase staining, red) were co-expressed in FRT control (A) or *psid-D4* MARCM clones (B-C) with or without co-expressed Psid rescue construct. *psid-D4* mutant clones (B) show similar levels of β-gal but decreased level of N-Drk-GFP in comparison to FRT control (A). (C) Expression of WT Psid rescued the level of N-Drk-GFP in *psid-D4* mutant clones. (D) Levels of N-Drk-GFP or WT GFP, normalized with β-gal level, in MARCM clones with indicated genotype were quantified and the average and standard deviation were shown (N≥6 for each genotype; * indicates P<0.0001 and x indicates P>0.05, t test). Data were collected from eye discs containing MARCM clones similar to those shown in (A-C). (E-J) Wing discs containing FRT control or *psid* MARCM clones marked by GFP expression. POE/Ubr4 RNAi clones were generated with the expression of p35 to prevent cell death. p35 expression alone did not influence Drk level (E). Ubr4/POE RNAi increased the Drk level in FRT control clones (F). In contrast, Ubr1 RNAi (G) or Ubr5 RNAi (H) clones did not affect Drk levels. Ubr4/POE RNAi restored the Drk level in *psid-D1* clones (I) but did not influence the level of PP2AC (J), suggesting a specific effect of Ubr4 on Drk but not PP2AC degradation.

Furthermore, the reduction in the level of GFP can be rescued by expressing Psid^WT^ and Psid^SA^ but not the NAA20 binding defective Psid^SD^ (Fig. 5A-D), similar to the observed effects of Psid on endogenous Drk protein levels as shown in Fig. 4C-F’. These results show that NatB regulates Drk protein stability through its N-terminal sequences.

### Drk is degraded by the N-end rule E3 ubiquitin ligase Poe/UBR4

The N-end rule pathway regulates protein stability through the N-terminal residues, which are often modified and recognized by the N-recognins [3]. Of the four N-recognins found in mammalian cells, only three (Ubr1, Ubr4, and Ubr5) are present in *Drosophila*. We tested the effect of knockdown fly Ubr1, Ubr4, or Ubr5 on Drk levels. As cells with Ubr4/Poe knockdown were mostly eliminated in developing discs, baculovirus p35 was expressed together with Ubr4 RNAi to inhibit cell death. Ubr4 knockdown with p35 expression significantly increased Drk levels in WT background (Fig. 5F-F’), while p35 expression alone did not significantly affect Drk levels (Fig. 5E-E’’). Furthermore, Ubr4 knockdown with p35 expression restored Drk levels in *psid* mutant clones (Fig. 5I-I’, compare with 4B-C’). On the other hand, knockdown Ubr1 or Ubr5 did not affect Drk levels (Fig. 5G-H’) and knockdown Ubr1 did not affect Drk levels in *psid* mutant background either (Fig. S4B-C’). Therefore, Drk protein degradation is regulated by the N-end rule E3 ubiquitin ligase Ubr4. These results, in conjunction with the finding that NatB regulates Drk stability through its N-terminal sequences, suggest that NatB regulates Drk protein stability through modification of its N-terminus, which alters its recognition by the N-end rule pathway.

### NatB decreases the levels of PP2AC, which inhibits MAPK activation in developing imaginal discs

While PP2A can potentially dephosphorylate multiple components of the Ras/MAPK pathway and exerts both negative and positive regulations, knockdown of PP2A subunits in *Drosophila* cells enhanced insulin-induced MAPK activation and reducing the gene dosage of PP2AC, the catalytic subunit of PP2A, stimulates activated Ras induced signaling in eye tissues [28, 29]. These observations suggest that PP2A has an overall negative effect on RTK/Ras induced MAPK activation *in vivo*.

We first used PP2AC RNAi to identify an antibody that can recognize the endogenous fly protein. As shown in Fig. 6E, PP2AC RNAi significantly decreased PP2AC protein signal, indicating that this PP2AC antibody can specifically detect the endogenous protein. Interestingly, MARCM clones with either *psid^D1^* or *psid^D4^* mutation showed increased PP2AC protein levels (Fig. 6A-B’). The increased PP2AC protein was dependent on the *psid* mutation since expression of WT Psid blocked the increased PP2AC levels (Fig. 6C-C’). Furthermore, knockdown of NAA20 also significantly increased PP2AC levels (Fig. 6D-D’). These results showed that inactivation of NatB activity increased the levels of PP2AC, the catalytic subunit of PP2A that negatively regulates EGFR/Ras-induced MAPK activation.

**Fig. 6.**
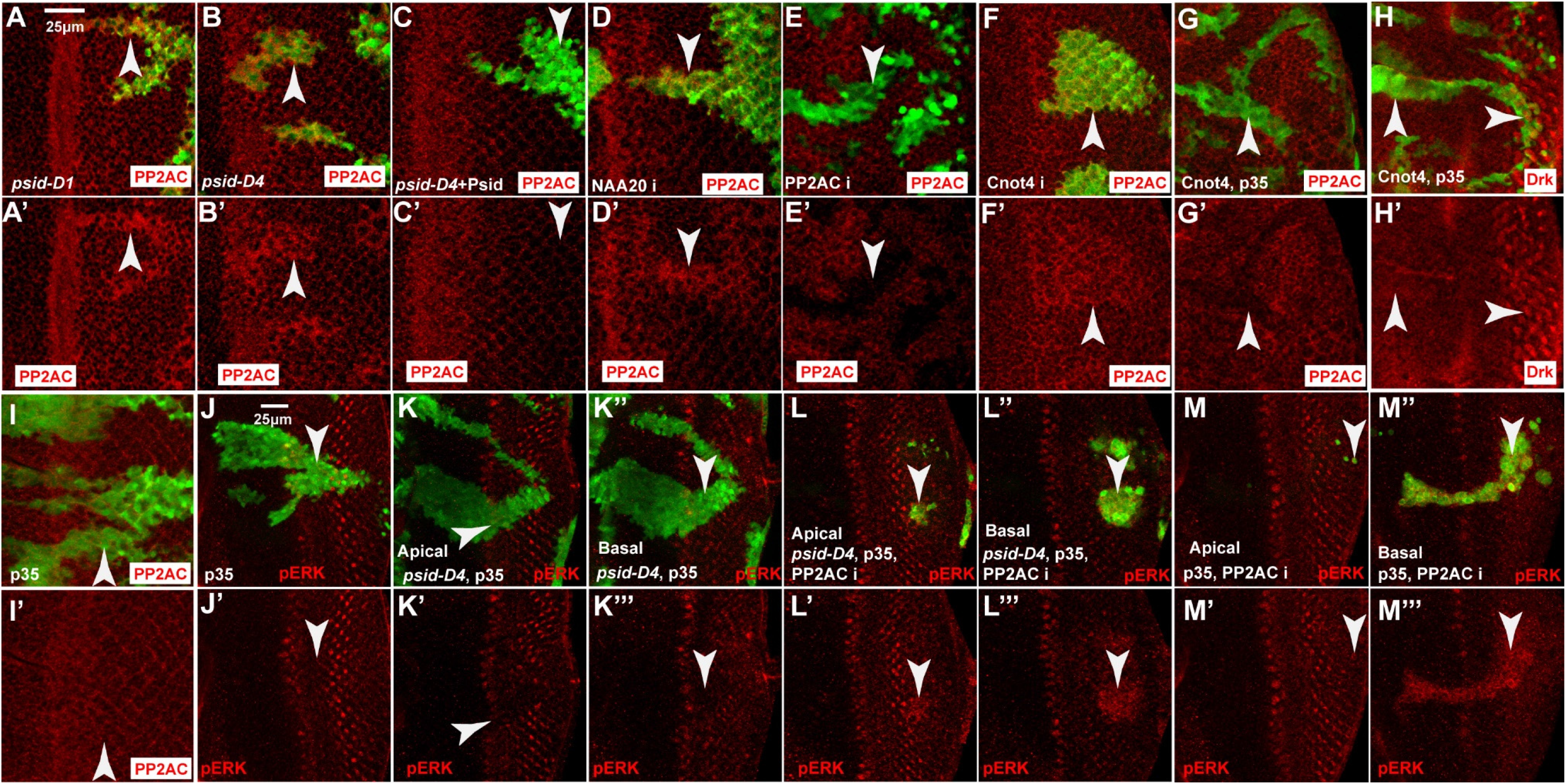
Inactivation of NatB increased levels of PP2AC, which contributes to the inhibition of MAPK activation by *psid* mutation. (A-C) Increased PP2AC level (anti-PP2AC staining, red) was observed in both *psid-D1* (A) and *psid-D4* (B) MARCM clones (marked by GFP expression) in eye discs. Expression of WT Psid (C) reduced PP2AC levels in *psid-D4* MARCM clones to background levels. (D) NAA20 RNAi clone generated with eyeflp-CoinGal4 and marked by GFP expression showed increased PP2AC level. (E) PP2AC RNAi expression in FRT82B MARCM clones decreased PP2AC levels in eye disc. (F) Cnot4 RNAi clone generated with eyeflp-CoinGal4 and marked by GFP expression showed increased PP2AC level. (G-I) GFP and p35 were expressed together with Cnot4 to prevent cell death and to mark the expressing cells. Cnot4 overexpression decreased PP2AC protein level (G) but did not significantly affect Drk level (H), while expressing p35 alone did not affect PP2AC level (I). (J-M) Apical or Basal sections of representative eye discs containing *psid* or FRT control MARCM clones marked by GFP expression. p35, which did not affect pERK (J), was used to prevent the elimination of PP2AC RNAi clones. *psid-D4* clones showed decreased pERK level (K). PP2AC RNAi expression cells in *psid-D4* clones (L-L”’) or in FRT control clones (M-M”’) were mostly located at the basal section (L”-L”’ and M”-M”’) and showed increased pERK levels.

Overexpression of PP2AC in mouse heart was shown to increase PP2AC level, reduce the phosphorylation level of its targets, and impair cardiac function [30]. To determine whether increased PP2AC levels contribute to reduced MAPK activation in *psid* mutant clones, we tested effects of knockdown PP2AC in *psid* mutant clones. Since PP2AC knockdown induced significantly levels of cell death, baculovirus p35 protein, which does not affect pERK levels when expressed alone (Fig. 6J), was expressed in conjunction with PP2AC knockdown. PP2AC-RNAi cells, which were observed mostly in the basal region of the eye disc even in the presence of p35 expression, showed increased pERK levels (Fig. 6M-M”’). This result is consistent with the previous finding that PP2AC has an overall negative effect on RTK/Ras induced MAPK activation [28]. Importantly, *psid* mutant cells with PP2AC knockdown showed increased pERK levels, in contrast to the reduced pERK levels observed in *psid* mutant clones (Fig. 6K-L”’). These results suggested that increased PP2AC contributes to the inhibition of MAPK activation by *psid* mutation.

### PP2AC and Drk are regulated by distinct branches of the N-end rule pathways

Since PP2AC levels are increased when NatB activity was inactivated, we hypothesized that the N-terminally acetylated PP2AC but not the N-terminally unmodified PP2AC is preferentially degraded. N-terminally acetylated proteins are degraded by the Ac/N-end rule pathway mediated by Doa10 or Not4 E3 ubiquitin ligases in yeast and the Doa10 homolog Teb4 in mammalian system [5]. The Doa10/Teb4 and Not4 homologs in *Drosophila* are CG1317 and Cnot4, respectively. We generated an allele of fly Teb4 that deleted a significant portion of the open reading frame including the initiation ATG (Fig. S5A). Clones of cells with *teb4/CG1317* mutation were quite small and did not affect PP2AC levels (Fig. S5B-B’). These results suggest that Teb4/CG1317 does not significantly affect PP2AC degradation. On the other hand, inactivating the fly Not4 E3 ubiquitin ligases homolog Cnot4 with RNAi significantly increased levels of PP2AC protein (Fig. 6F-F’) but not PP2AC mRNA (Fig. S5C). Additionally, overexpression of Cnot4 significantly decreased the basal levels of PP2AC (Fig. 6G-G’). These results suggest that Cnot4 but not Teb4/CG1317 contribute to the degradation of PP2AC.

The above results showed that PP2AC is degraded by the Ac/N-end rule E3 ubiquitin ligase Cnot4 while Drk is degraded by the Arg/N-end rule pathway E3 ubiquitin ligase Ubr4. We further determined whether PP2AC could also be significantly affected by the Arg/N-end rule pathway and whether Drk significantly affected by the Ac/N-end pathway. Ubr4 knockdown did not significantly affect PP2AC levels (Fig. 5J-J’) despite significantly increased the levels of Drk (Fig. 5F-F’). In addition, knockdown of Ubr1 and Ubr5, the other two fly N-recognins of the Arg/N-end rule pathway, did not significantly affect PP2AC levels either (Fig. S5D-E’). These results suggest that PP2AC is not significantly affected by the Arg/N-end rule pathway E3 ubiquitin ligases. On the other hand, overexpression of Cnot4 did not significantly affect Drk levels despite significantly reduced PP2AC levels (Fig. 6G-H’). In addition, mutation of Teb4/CG1317 did not significantly affect Drk levels either (Fig. S5F). These results show that Drk protein level was not significantly affected by the Ac/N-end rule E3 ubiquitin ligases.

Taken together, our results show that PP2AC is preferentially targeted for degradation by the Ac/N-end rule pathway E3 ubiquitin ligase Cnot4 while Drk is preferentially targeted for degradation by the Arg/N-end rule pathway E3 ubiquitin ligase Ubr4.

### NatB inactivation decreased the level of MAPK, another positive component of the EGFR/MAPK pathway

The above results showed that NatB inactivation inhibited EGFR/MAPK pathway by decreasing the levels of the conserved positive regulator Drk and increasing the levels of the conserved negative regulator PP2AC. To gain better understanding of how NatB/N-end rule pathway modulate EGFR/MAPK signaling, we characterized the effect of NatB on two additional potential targets, Spry and MAPK. MAPK has high level of overall sequence conservation except the N-terminal region. Interestingly, a previous study showed that MAPK is degraded by the Poe/Ubr4 E3 ubiquitin ligase, suggesting that MAPK is regulated by the Arg/N end rule pathway [31]. Therefore, it is very interesting to determine whether NatB affects MAPK levels.

We found that inactivation of NatB with *psid* mutation moderately reduced MAPK levels in wing disc (Fig. 7C), using an anti-MAPK antibody that can detect the endogenous MAPK protein (Fig. 7F). In addition, significant reduced MAPK protein levels were also observed in both *psid^D1^* or *psid^D4^* mutant clones in fat body (Fig. 7A-B’). Therefore, NatB also regulates MAPK levels. In support of the previous report that MAPK is degraded by Ubr4, knockdown of Ubr4 with p35 expression significantly increased MAPK levels (Fig. 7D-D’), while p35 expression alone or knockdown of Ubr1 or Ubr5 had no effect (Fig. 7G-I’). In addition, knockdown of Ubr4 also blocked *psid* mutation induced-decrease in MAPK levels (Fig. 7C-E’). Taken together, NatB activity also increase the level of MAPK, another positive component of the EGFR signaling pathway.

**Fig. 7.**
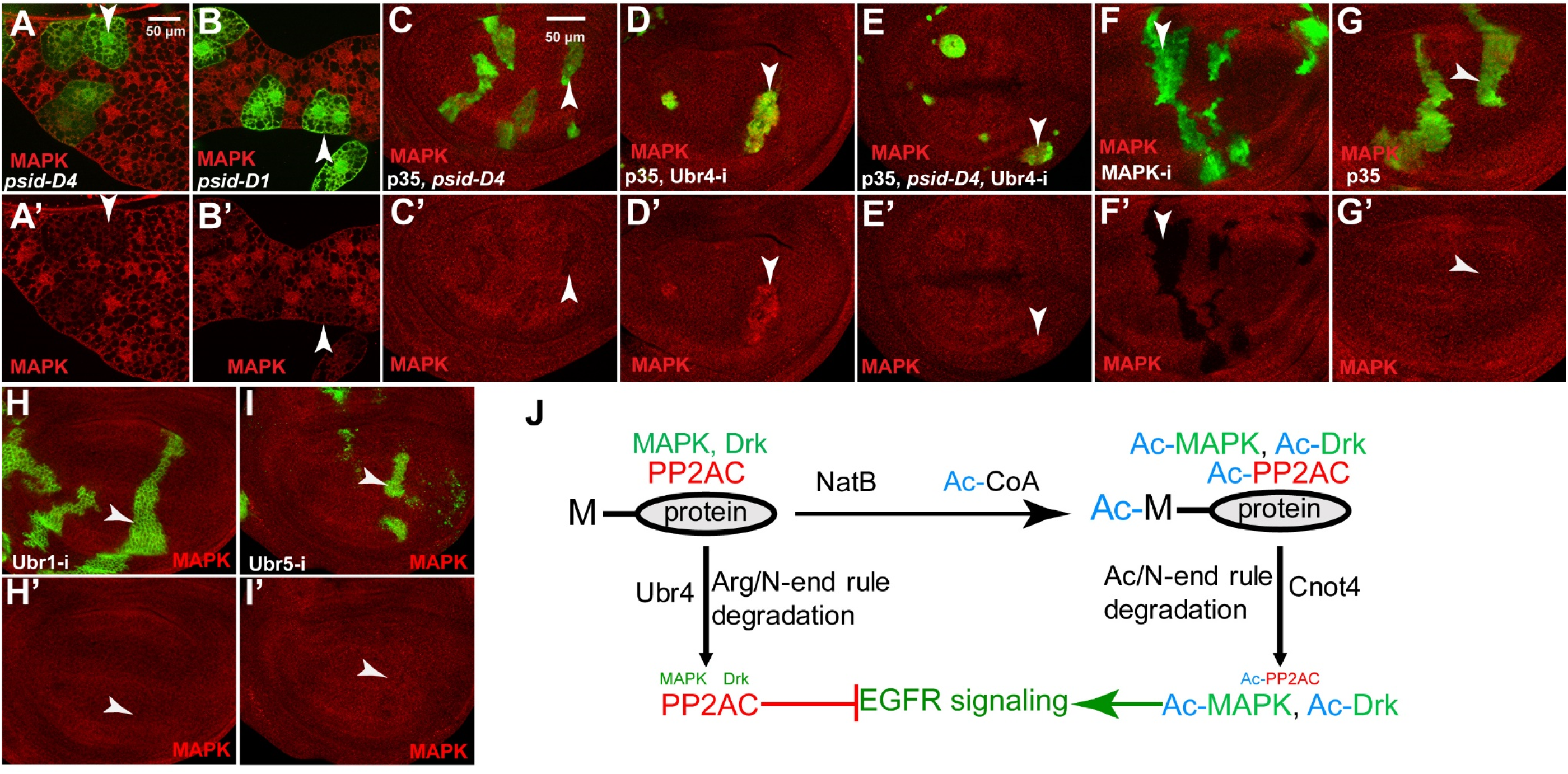
NatB inactivation decreased the level of MAPK and proposed model of EGFR/MAPK signaling regulation by NatB and the N-end rule pathways. (A-B) *psid-D1* (A) or *psid-D4* (B) clones in 3rd instar fat body, which were marked by GFP expression, showed decreased MAPK levels (anti-MAPK staining, red). (C) *psid-D4* clones in wing disc, marked by GFP expression, showed decreased MAPK level. (D-E) Ubr4 RNAi clones, marked by GFP expression, showed increased MAPK level (D). Ubr4 RNAi also restored the level of MAPK in *psid-D4* clones (C, E). p35 expression, which did not affect MAPK levels (G), was used to block the death of Ubr4 RNAi cells. (F) The specificity of the anti-MAPK antibody. Reduced MAPK staining was observed in MAPK RNAi clones marked by GFP expression. (H-I) Ubr1 RNAi (H) or Ubr5 RNAi (I) clones did not significantly affect MAPK levels. (J) Proposed model of EGFR/MAPK signaling regulation by NatB and the N-end rule pathways through inversely modulating the levels of positive and negative components of the EGFR/MAPK pathway. Drk/Grb2 and MAPK, two positive components of the EGFR signaling pathway are targeted for degradation by the Arg/N-end rule pathway and E3 ubiquitin ligase Ubr4 while PP2AC, a negative regulator of EGFR signaling pathway, is targeted for degradation by the Ac/N-end rule pathway E3 ubiquitin ligase Cnot4. The level of acetylation of these proteins are controlled by NatB activity and the key rate-limiting substrate Ac-CoA. High NatB activity and high Ac-CoA level will increase the levels of Nt-acetylated Grb2/Drk, MAPK, and PP2AC, leading to high EGFR signaling due to elevated levels of Grb2/Drk and MAPK (positive components of the pathway) and reduced levels of PP2AC (negative component of the pathway). In contrast, low NatB activity and low Ac-CoA levels will reduce the levels of Nt-acetylated Grb2/Drk, MAPK, and PP2AC, leading to reduced EGFR signaling due to decreased levels of Grb2/Drk and MAPK (positive components of the pathway) and increased levels of PP2AC (negative component of the pathway).

In contrast to the highly conserved sequences of Drk and PP2AC, Spry has low overall sequence conservation except in a cysteine-rich region near the C-terminus [32]. Staining with an antibody against Spry [32] detected elevated signals in posterior eye discs that was significantly reduced by Spry RNAi (Fig. S6A-A’, white arrowheads) and background signals in other parts of eye discs that was not affected by Spry-RNAi (Fig. S6A-A’, yellow arrowheads). This is consistent with the observations that Spry expression is dependent on RTK signaling [32, 33]. Decreased levels of Spry was detected in *psid* mutant clones (Fig. S6B). Since *psid* mutation reduces EGFR/MAPK signaling, which in turn regulates Spry expression, the observed decreased Spry level in posterior eye disc could be due to reduced EGFR signaling in *psid* clones. As we could not detect basal levels of Spry in other parts of discs with no RTK signaling, we could not exclude the possibility that NatB may affect Spry protein stability in addition to the role on Spry expression.

## Discussion

Our results suggest that NatB activity modulates EGFR signaling by altering the levels of multiple positive and negative components of the pathway. The N-terminus of protein have been shown to influence protein stability through the N-end rule pathway with branches that selectively degrade N-terminally acetylated protein or N-terminally unmodified protein [5]. Interestingly, we found that NatB targets that are highly conserved positive components of EGFR signaling pathway are selectively targeted for degradation by the Arg/N end rule E3 ubiquitin ligase Ubr4 (Fig. 7J). In contrast, NatB target that is a highly conserved negative component of the pathway, PP2AC, is selectively targeted for degradation by the Ac/N end rule E3 ubiquitin ligase Cnot4 (Fig. 7J). As E3 ubiquitin ligases for the two branches of the N-end rule pathway selectively recognize the positively charged non-acetylated N-terminus or the acetylated N-terminus that lacks the positive charge, respectively, NatB activity, which shifts the level of protein Nt-acetylation, will lead to coordinated changes in the levels of positive and negative components of the pathway (Fig. 7J, S7A-B). Therefore, inhibition of NatB activity will results in the loss of N-terminal acetylated Drk/Grb2 and MAPK and the degradation of the N-terminal non-acetylated proteins, leading to decreased levels of positive components (Fig. S7A). Furthermore, inhibition of NatB activity will also results in accumulation of N-terminal non-acetylated PP2AC and prevents its degradation by Ac/N end rule E3 ubiquitin ligase Cnot4, leading to increased levels of the negative component of the pathway (Fig. S7A). The inverse change in the levels of positive and negative components of the pathway underlie NatB activity-mediated regulation of EGFR signaling (Fig. 7J, S7A-B).

Although N-terminal acetylation was generally viewed as a constitutive, co-translational modification that does not have regulatory function, our results suggest N-terminal acetylation by NatB may provide important regulatory function. First, our results show that NatB activity coordinately regulates the levels of positive and negative components of the EGFR signaling pathway. Second, we found that knockdown of Ubr4 in WT background induced significant elevated levels of both Drk/Grb2 and MAPK. As Ubr box proteins specifically recognizes proteins with non-acetylated proteins, these results suggest that significant levels of Drk/Grb2 and MAPK are synthesized without N-terminal acetylation in normal growth conditions and these proteins are targeted for degradation by Ubr4. It is likely that these proteins are partially Nt-acetylation under normal growth conditions, which could potentially be regulated by increasing or decreasing the level of NatB activity or Acetyl-CoA, a key substrate of N-terminal acetylation.

Acetyl-CoA has been suggested as a central metabolite and second messenger of the cellular energetic state [34]. The levels of Acetyl-CoA, which is influenced by the availability of nutrients and growth factors, can vary significantly in cells and influence the levels of protein acetylation by various acetyl transferases [34, 35]. Indeed, Acetyl-CoA levels was shown to regulate the N-terminal acetylation of several NatA targets, which modulates the sensitivity of cells to apoptosis [36]. As the Km for NatB from *Candida* is quite high (around 50 µM) [37], it is likely that N-terminal acetylation by NatB will also be influenced by Acetyl-CoA levels, which will link the regulation of EGFR signaling with cellular energetic status. Therefore, our study suggest a novel mechanism by which the activity of EGFR signaling is regulated by the cellular energetic status.

In addition to Acetyl-CoA level, NatB activity may also be regulated. It was shown that a conserved serine in Psid just downstream of the NAA20 interaction domain regulated NatB complex formation in a phosphorylation-dependent manner (Stephan et al., 2012). Similarly, we found that expressing the phosphorylation resistant but not the phosphorylation mimic form of Psid could rescue the effect of *psid* mutation on EGFR signaling. These results suggest NatB activity is regulated by cellular signaling pathways through phosphorylation. Taken together, our study suggest that the level of N-terminal acetylation is potentially regulated by cellular signaling and nutrient status, which can in turn regulate the strength of EGFR signaling output by coordinately regulated the levels of multiple positive and negative regulators of the pathway (Fig. S7A-B). The high sequence conservation of these EGFR signaling components suggest that the mechanisms we uncovered in *Drosophila* will likely be conserved in mammalian systems. Indeed, a recent study found that inhibition of NatB significantly decreased EGF-induced ERK activation in human liver cancer cells [38]. As mutation or overexpression of EGFR or Ras that deregulate EGFR signaling are quite common in human cancers and NatB subunits are significant unfavorable prognostic markers for human cancers [39-41], our results could potentially provide a new strategy to develop therapeutic interventions for these cancers.

The regulation of MAPK activity by NatB described in this study also provides a mechanism by which Nt-acetylation by NatB is required to ensure olfactory receptor neuron (ORN) survival as described previously [20]. It was shown that the phospho-resistant Psid^SA^ mutant rescued the *psid* cell-loss phenotype while the phosphomimetic Psid^SD^ mutant failed to rescue [20]. In this study, we showed that the phospho-resistant Psid^SA^ but not the phosphomimetic Psid^SD^ was able to rescue *psid* mutation-mediated MAPK inhibition and *rbf psid* synergistic cell death.

We showed that *psid* mutation significantly inhibited dpERK level induced by activated EGFR and Ras but not activated Raf. At first glance, these results do not seem to be consistent with the result that MAPK is also regulated by NatB. The most likely explanation is that the effect of reduced MAPK levels is compensated by the effect of increased levels of PP2AC. It was shown that activation of Raf by Ras-GTP involve activating phosphorylation as well as inhibitory phosphorylation by ERK feedback regulation [42]. It is possible that in experiments involving activated Raf-induced MAPK activation, the main function of PP2A is to remove the inhibitory phosphorylation on Raf. In contrast, in experiments involving activated Ras-induced MAPK activation, the dominant function of PP2A is to remove the activating phosphorylation. Consistent with this, reducing the PP2AC gene dosage was found to impair signaling from activated Raf but stimulate signaling from activated Ras [28]. Therefore, *psid* mutation induced increased PP2AC potentially compensated decreased MAPK levels, resulting no obvious inhibition of activated Raf-induced dpERK level.

## Methods

### Drosophila stocks and genetics

The following fly stocks were used in this study: *rbf^15aΔ^* [1^7^]; *psd-D1*(BL41122, BL indicating Bloomington Drosophila stock center); *psd-D4* (BL41123), UAS-Rbf RNAi (BL36744), *scrib* ^673^ (BL41175), UAS-NAA20 RNAi (BL36899), UAS-Drk RNAi (BL41692), Drk-PZ(BL12378), UAS-Poe RNAi (BL32945), UAS-PP2AC RNAi (BL 27723), UAS-Cnot4 RNAi (BL42513), UAS-Cnot4 (BL22246), UAS-p35 (BL5072, BL5073), UAS-MAPK RNAi (BL34855), UAS-Ubr5 RNAi (BL32352), UAS-Ubr1 RNAi (BL31374), UAS Psid-WT [19], UAS-Psid-S678D and UAS-Psid-S678A [20], aos-lacz and UAS-EGFR*^CA^* [1^3^], UAS-ras*^v12^* (BL64196), UAS-raf*^gof^* (BL2033), UAS-GFP (BL5431), CoinFLP-Gal4-UAS-GFP (BL58751), UAS-Dcr2 (BL58757), UAS-sprouty RNAi (BL36709), and UAS-sprouty [33].

Main genetic technologies used in this study include: FLP/FRT system to generate regular loss of function mosaic clones [43]; MARCM system to generate mosaic clones with both mutation and ectopic expression [44]; UAS/Gal4 and Flp-out or CoinFLP system to induce ectopic expression of RNAi in clones or in whole eyes [45-47].

### Genetic screen for *rbf* synthetic lethal mutations and generation of N-Drk-GFP transgenic fly and CG1317/Teb4 deletion allele

Ethyl methanesulfonate (EMS)-induced mutant screen was carried out as described [17]. Isogenized w; *P{ry+, neoFRT82B}* males were used for mutagenesis, *rbf^15aΔ^,w, eyFLP*; *P{ry+, neoFRT82B} P{w+, Ubi-GFP} P{w+, Rbf-G3}* and *w, eyFLP*; *P{ry+,neoFRT82B} P{w+, Ubi-GFP}* stocks were used for screening and *rbf* dependence test.

The N-Drk-GFP transgenic fly, which contains the N-terminal 10 amino acid sequence from Drk (ATG GAA GCG ATT GCC AAA CAC GAT TTC TCT) fused to the N-terminus of GFP from PX458 plasmid was generated by PCR and verified by sequencing. The N-Drk-GFP fusion were cloned into the pUAST plasmid and transgenic flies were established.

The CG1317/Teb4 deletion allele was generated by crossing the P-element insertion line (BL20646) with the transposase line Δ2-3. Independent excision lines that have lost eye color were established. Deletions were identified by PCR using P element primers and primers flanking the P element. The breakpoints were determined by sequencing of the PCR products.

### Immunostaining, BrdU incorporation, and antibodies

Immunostaining was performed as previously described [13]. For dpErk staining, dissected discs were fixed with 8% formaldehyde in PBS for 1 hr. For PP2AC staining, dissected discs were fixed with 4% formaldehyde in 100mM Lysine for 1 hr on ice, and saponin was added in the blocking solution to a final concentration of 0.2% during the process of blocking and primary antibody incubation. For Brdu incorporation of male gonads, male gonads were dissected from larvae with growing tumors at 7-8 day after egg laying in Schneider’s medium. Samples were incubated with BrdU (75 μg/ml in Schneider’s medium) at RT for 1 hr, washed with PBS, and fixed with 4% formaldehyde in PBS for 30 minutes at RT, followed by postfixing with 4% formaldehyde in PBS with 0.6% Tween 20 for 30 minutes at RT. These samples were washed with DNase I buffer, followed by incubation with DNase I (100 U/500 μl) for 1 hr and wash with PBST (0.3% triton X-100). Primary antibodies used in this study: rabbit anti-activated Caspase-3 (C3, 1:400 from Cell Signaling), rabbit anti dpErk (1:400, Cell signaling), rabbit anti-MAPK (1:500, Cell signaling), mouse anti-PP2AC (1:400, Santa Cruz Biotechnology), rabbit anti-Drk (1:1000) [27], Guinea pig anti-senseless (1:1000, gift from Dr. Hugo Bellen), rat anti-Elav (1:100, DSHB), mouse anti-BrdU (1:50, DSHB), mouse anti-β-Galactosidase (1:100, DSHB). Secondary antibodies are from Jackson ImmunoResearch (1:400). Samples were mounted in 70% Glycerol with 1,4-diazabicyclo[2.2.2]octane (DABCO) at 12.5 mg/mL. Samples were imaged with an AxioCam CCD camera mounted on a Zeiss Axio Imager with ApoTome using the Zeiss Axiovision software.

### Quantification of cell death levels and N-Drk-GFP levels in developing imaginal discs

Cell death level was determined by the percentage of clone area (pixels) that have above background levels of caspase 3 (C3) signal using the Histogram function in Photoshop as described previously [21]. Background level of C3 signal was determined from the adjacent WT tissues that have no apoptosis. The average and standard deviation of percent cell death for each genotype discs was then determined from at least six imaginal discs and then compared. Two-way student T test was used to determine the significance of statistical differences between different genotypes.

To investigate the effect of *psidin* mutation on N-Drk-GFP levels, MARCM clones were generated and marked by LacZ expression. LacZ expression levels were determined by anti-β-gal staining and used as an internal control. Exposure time of imaging was optimized with the brightest samples and used for all samples. GFP and β-gal signal brightness were calculated using the Histogram function in Photoshop. Normalized GFP levels were showed as relative ratios of signal brightness of GFP to that of β-gal with the normalized GFP level in WT control set as 1. The average and standard deviation of relative GFP levels for each genotype discs was then determined using at least six imaginal discs and compared. Two-way student T test was used to determine the significance of statistical differences between different genotypes.

### RNA isolation and quantitative real-time PCR

Total RNA was extracted using TRI reagent (Invitrogen) from about 30 eye/antenna discs dissected from 3rd instar larvae cultured at 25 °C with expression of RNAi constructs driven by eyFLP, Act>CD2>Gal4. cDNA synthesis and qRT-PCR reactions were performed as previously described [48]. Ribosomal protein gene rp49 was used as an internal control in qPCR analysis. The averages and standard deviations of at least three independent replicates were shown. Primers used were as follows: PP2AC for 5’-GCAATCAGTTGACAGAGACACA-3’; PP2AC Rev 5’-CACCGGGCATTTTACCTCCT-3’; Rp49 F 5’-ACAGGCCCAAGATCGTGAAGA-3’; Rp49 R 5’-CGCACTCTGTTGTCGATACCCT-3’.

### Genotype of flies used in this study

Fig. 1

~~~
w, eyFLP /Y; FRT82B, Ubi-GFP /FRT82B
*rbf^15aΔ^*,w, eyFLP /Y; FRT82B, RBF-G3, Ubi-GFP/FRT82B
w, eyFLP /Y; FRT82B, Ubi-GFP /FRT82B, *psid* (*AE*,*EQ, psdin D1 or D4*)
*rbf^15aΔ^*,w, eyFLP /Y; FRT82B, RBF-G3, Ubi-GFP / FRT82B, *psid* (*AE*,*EQ, psdin D1 or D4*)
eyFLP, Act>CD2>Gal4/Y; UAS-Rbf RNAi /+ or UAS-NAA20 RNAi
eyFLP, Act>CD2>Gal4/Y; + or UAS-NAA20 RNAi
w,eyFLP (or HsFLP)/Y; Act > y >Gal4, UAS-GFP; FRT82B, tub-Gal80/ FRT82B, *psid D4*
*rbf^15aΔ^*, w,eyFLP (or HsFLP)/Y; Act > y >Gal4, UAS-GFP; FRT82B, RBF-G3, tub-Gal80/ FRT82B or (FRT82B, *psidin D4*)
*rbf^15aΔ^*,w, eyFLP (or HsFLP)/Y; Act > y >Gal4, UAS-GFP/ UAS-psidin WT; FRT82B, RBF-G3, tub-Gal80/ FRT82B, *psid D4*
~~~

Fig. 2

~~~
w, eyFLP /Y; FRT82B, Ubi-GFP / aos-lacz, FRT82B, *psdin* (*AE*,*EQ, psid D1 or D4*)
HsFLP; Act > y >Gal4, UAS-GFP/ + or UAS-psidin (WT, SD, or SA); FRT82B, tub-Gal80/ FRT82B,*psid D4*
eyFLP, UAS-Dcr2 / +; CoinFLP-Gal4-UAS-GFP; aos-lacz, UAS-NAA20 RNAi
~~~

Fig. 3

~~~
HsFLP; Act > y >Gal4, UAS-GFP/ UAS-(EGFR^CA^, Ras^v12^, or Raf^gof^); FRT82B, tub-Gal80/ FRT82B or (FRT82B, *psid D4*)
HsFLP; Act > y >Gal4, UAS-GFP/ UAS-EGFR^CA^, UAS-psid (WT, SD, or SA); FRT82B, tub-Gal80/ FRT82B, *psid D4*
eyFLP, UAS-Dcr2 / +; CoinFLP-Gal4-UAS-GFP/ EGFR^CA^; + or UAS-NAA20 RNAi
*rbf^15aΔ^*,w, HsFLP/Y; Act > y >Gal4, UAS-GFP/ + or UAS-(EGFR^CA^, Ras^v12^, or Raf^gof^); FRT82B, RBF-G3, tub-Gal80/ FRT82B, *psid D4*
~~~

Fig. 4

~~~
w, HsFLP; FRT82B, Ubi-GFP / FRT82B, *psdin D1*
HsFLP; Act > y >Gal4, UAS-GFP/ + or UAS-psid (WT, SD, or SA); FRT82B, tub-Gal80/FRT82B, *psid D4*
eyFLP, UAS-Dcr2 / +; CoinFLP-Gal4-UAS-GFP; UAS-NAA20 RNAi
eyFLP, UAS-Dcr2 / +; CoinFLP-Gal4-UAS-GFP; UAS-Drk RNAi
eyFLP, UAS-Dcr2 / +; CoinFLP-Gal4-UAS-GFP/ UAS-EGFR^CA^; + or UAS-Drk RNAi
eyFLP, UAS-Dcr2 / +; CoinFLP-Gal4-UAS-GFP/ UAS-EGFR^CA^,UAS-sprouty; + or UAS-Drk RNAi
~~~

Fig. 5

~~~
eyFLP; Act > y >Gal4, UAS-Lacz; FRT82B, tub-Gal80/ UAS-N-Drk-GFP, FRT 82B
eyFLP; Act > y >Gal4, UAS-Lacz / + or UAS-psid (WT, SD, or SA); FRT82B, tub-Gal80/ UAS-N-Drk-GFP, FRT 82B, *psid D4* (*or psid D1*)
eyFLP; Act > y >Gal4, UAS-Lacz / Act > y >Gal4, UAS-GFP; FRT82B, tub-Gal80/ FRT 82B or (FRT82B, *psid D4*)
HsFLP; Act > y >Gal4, UAS-GFP/UAS-p35; FRT82B, tub-Gal80/ FRT82B
HsFLP; Act > y >Gal4, UAS-GFP/UAS-p35; UAS-Ubr4 RNAi,FRT82B, tub-Gal80/ FRT82B or (FRT82B, *psid D1*)
HsFLP; tub-Gal80, FRT40A/ FRT40A; Act > y >Gal4, UAS-GFP / UAS-(Ubr1 RNAi or Ubr5 RNAi)
~~~

Fig. 6

~~~
HsFLP; Act > y >Gal4, UAS-GFP; FRT82B, tub-Gal80/ FRT82B, *psid D4 or* (*psid D1*)
HsFLP; Act > y >Gal4, UAS-GFP/ UAS-psid WT; FRT82B, tub-Gal80/ FRT82B, *psid D4*
eyFLP, UAS-Dcr2 / +; CoinFLP-Gal4-UAS-GFP; UAS-NAA20 RNAi
w,eyFLP; Act > y >Gal4, UAS-GFP; FRT82B, tub-Gal80/UAS-PP2AC RNAi, FRT82B
eyFLP, UAS-Dcr2 / +; CoinFLP-Gal4-UAS-GFP/ UAS-Cnot4 RNAi
HsFLP; Act > y >Gal4, UAS-GFP, UAS-p35/ + or UAS-Cnot4; FRT82B, tub-Gal80/ FRT82B
HsFLP; Act > y >Gal4, UAS-GFP/ UAS-p35; FRT82B, tub-Gal80/ FRT82B or ((FRT82B, *psid D4*), (UAS-PP2AC RNAi, FRT82B, *psid D4*) or (UAS-PP2AC RNAi, FRT82B))
~~~

Fig. 7

~~~
HsFLP; Act > y >Gal4, UAS-GFP; FRT82B, tub-Gal80/ FRT82B, *psid D4 or* (*psid D1*)
HsFLP; Act > y >Gal4, UAS-GFP/UAS-p35; FRT82B, tub-Gal80/ FRT82B or (FRT82B, *psid D4*)
HsFLP; Act > y >Gal4, UAS-GFP/UAS-p35; UAS-Ubr4 RNAi,FRT82B, tub-Gal80/ FRT82B or (FRT82B, *psid D4*)
HsFLP; tub-Gal80, FRT40A/ FRT40A; Act > y >Gal4, UAS-GFP / UAS-MAPK RNAi or Ubr5 RNAi
HsFLP, Act>CD2>Gal4/Y; UAS-Ubr1 RNAi
~~~

## Acknowledgments

We thank Tianyi Zhang, Xun Pei, Yang Liao, Zhenyu Zhang, Xuan Li and Jiehui Zhang from the Du lab for assistance in *rbf*-dependent genetic screening and mapping, fly stock maintenance, and plasmid construction. We would like to thank Drs. Efthimios Skoulakis, Denise Montell, Ilona Grunwald Kadow, Hugo Bellen, Matthew Freeman and Mark Krasnow for providing fly stocks and antibodies. We thank the Bloomington Stock Center (NIH P40OD018537) for providing fly stocks and the Developmental Studies Hybridoma Bank (DSHB, created by the NICHD of the NIH and maintained at The University of Iowa) for providing antibodies. This work is supported by a grant from National Institute of Health R01 GM120046.

## Declaration of Interests

The authors declare that they have no competing interest with the contents of this article.

## Author contributions

ZS and WD designed the study, interpreted the results, and wrote the manuscript. ZS carried out experiments and collected data. Both authors read and approved the final manuscript.

