## supplemental figures for "NatB modulates Rb mutant cell death and tumor growth by regulating EGFR/MAPK signaling through the N-end rule pathways"

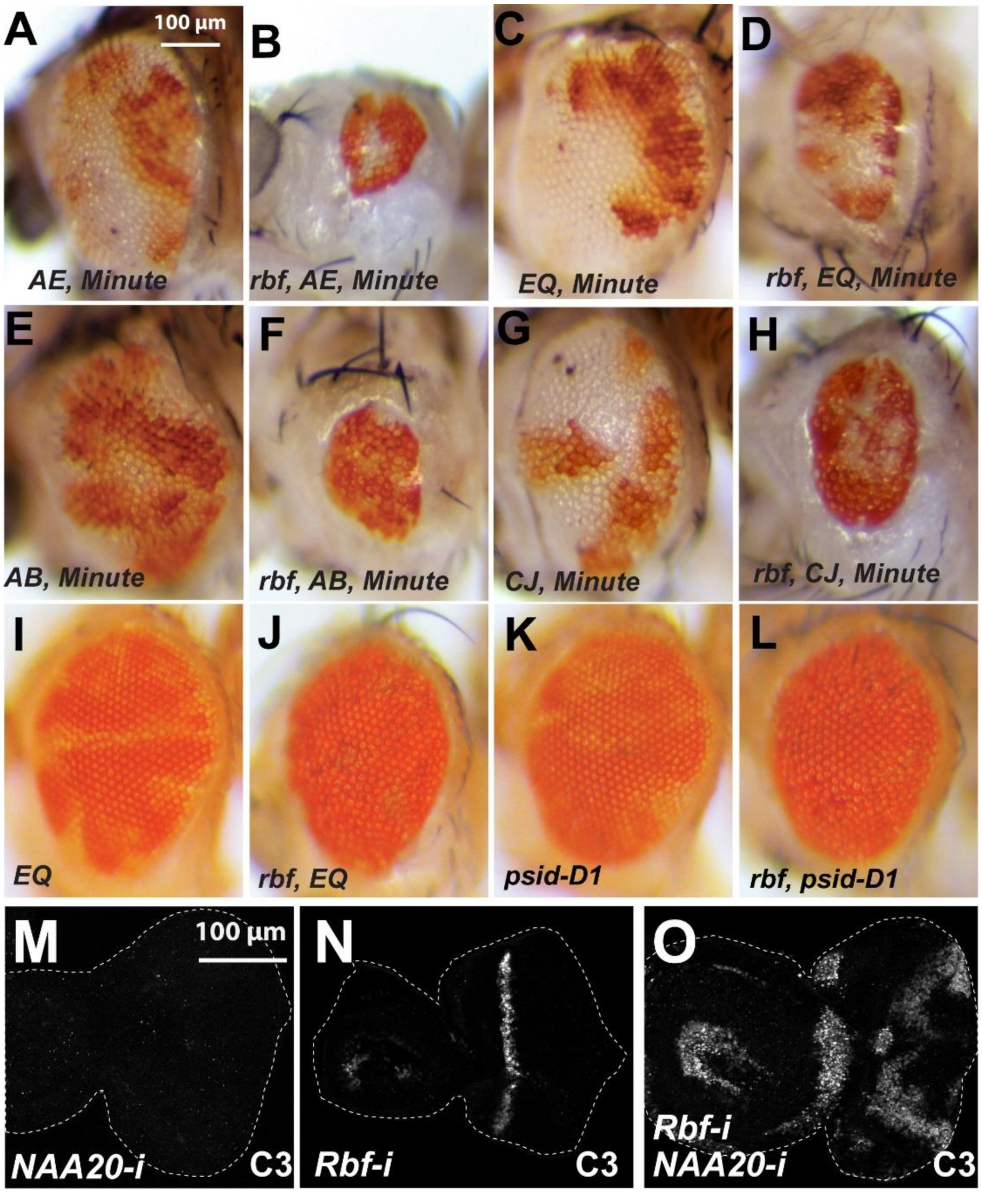


**Fig. S1. Inactivation of *rbf* and NatB subunits induce loss of double mutant tissues and synergistic cell death.** (A-H) Large clones (white patches) of *psidin* alleles (A,C,E and G) were observed when mutant clones were generated in a *Minute* background. In contrast, *rbf psidin* double mutant clones generated in the *Minute* background were mostly lost, resulting in smaller eyes (B,D,F and H). (I-L) Adult eyes with single mutant clones of additional *psidin* alleles or double mutant clones of *psidin*, *rbf*. Mutant clones were marked by the lack of red pigment. (M-O) Cell death level (caspase 3 staining) in third instar eye/antenna discs with single or double RNAi of Rbf or NAA20.

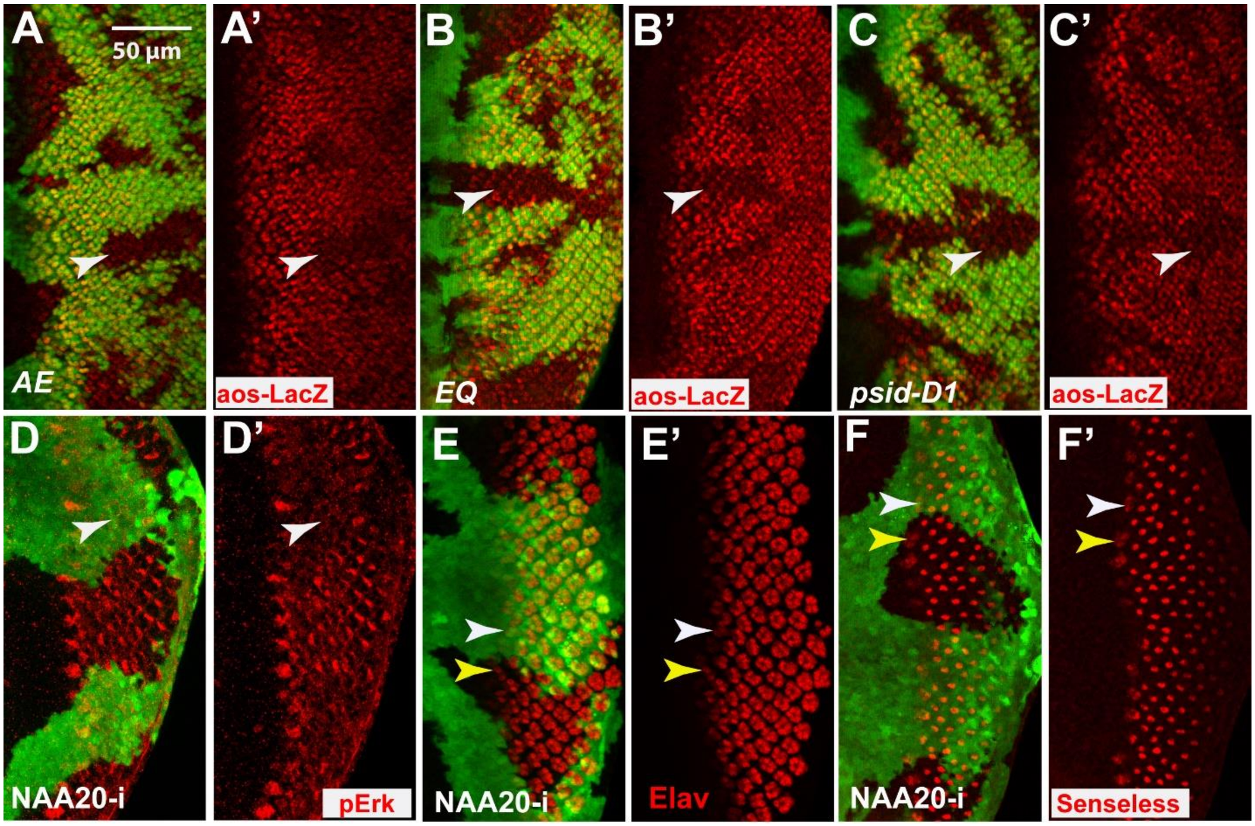


**Fig. S2. Inactivation of NatB subunits reduced EGFR signaling and slightly delayed neuronal differentiation.** (A-C') Expression levels of aos-lacZ was reduced in mutant clones of additional *psidin* alleles marked by the absence of GFP. (D-F') NAA20 RNAi clones generated with EyeFlp-CoinGal4 and marked by GFP expression were stained with pErk (D-D'), and neuronal differentiation markers Elav (E and E') and Senseless (F and F').

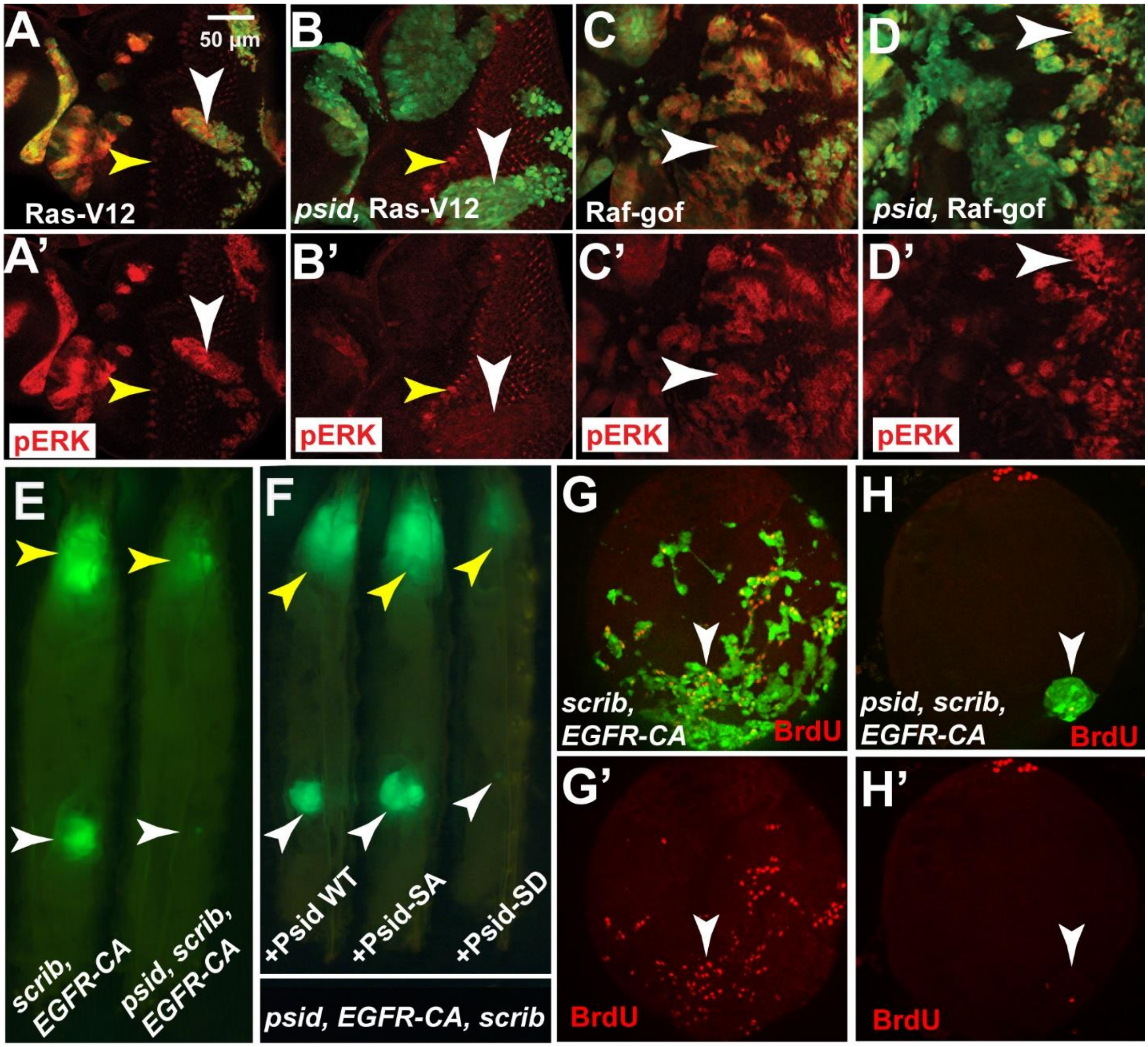


**Fig. S3. Effect of *psid* mutation on activated Ras and Raf-induced MAPK activation in eye discs and on activated EGFR signaling induced tumor growth.** (A-D) The effects of *psid-D4* mutation on activated Ras (A-B) or activated Raf (C-D) induced MAPK activation in eye discs were detected by pERK staining (red). Activated Ras-induced pERK (white arrowhead in A) was much higher than the endogenous pERK observed in the morphogenetic furrow (Yellow arrowhead in A). *psid-D4* mutation significantly reduced activated Ras-induced pERK (white arrowhead pointed in B), which was similar to the endogenous pERK level in WT tissues (yellow arrow pointed in B). Activated Raf-induced pERK (white arrow pointed in C) was not obviously affected by *psid-D4* mutation (white arrow pointed in D). (E) *psid* mutation inhibited Ey-FLP induced MARCM clones of activated EGFR expression (labelled with GFP) in *scrib* mutant clones. Yellow and white arrows point to tumor growth in the head region and the male gonads, respectively. (F) Expression of WT Psid or the Psid-SA mutant but not the NAA20-binding defective Psid-SD mutant restored tumor growth, suggesting the NatB activity is required for activated EGFR-induced tumor growth. (G-H) *psid* mutation significantly inhibited activated EGFR-induced cell proliferation in male gonads. Activated EGFR expression in *scrib* mutant clones were highly proliferative with high level of BrdU incorporation (red). In contrast, mutation of *psid* significantly inhibited BrdU incorporation in *EGFR scrib* cells.

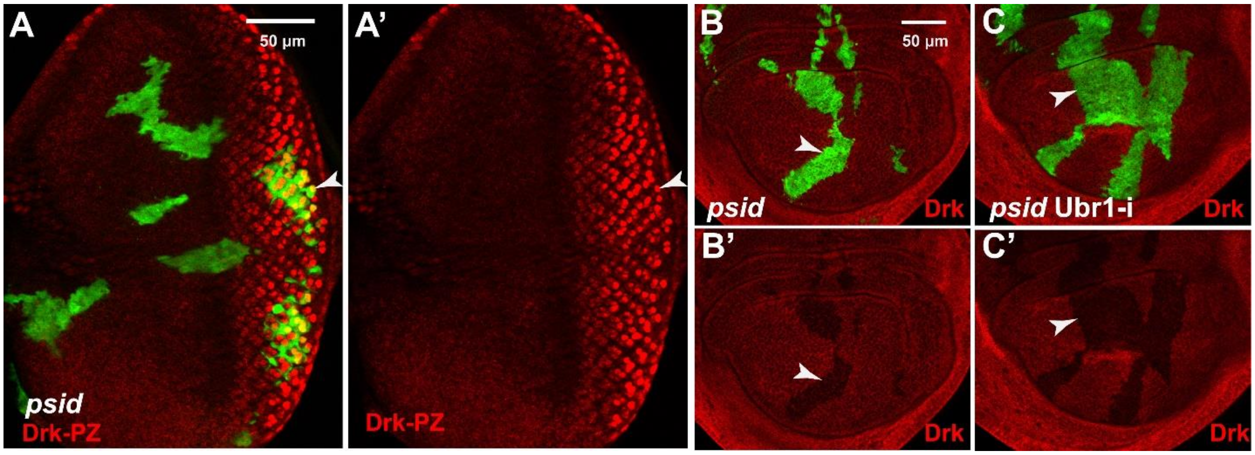


**Fig. S4**. **Inactivation of Psid did not reduce the Drk reporter expression and Knockdown E3 ubiquitin ligase Ubr1 did not restore Drk protein level in *psid* mutant clones**.

(A) *psid-D1* MARCM clones in eye discs, marked by GFP expression, did not significantly affect Drk expression as shown by level of β-gal (red) expression from a Drk enhancer trap line. (B) *psid-D1* MARCM clones in wing discs, marked by GFP expression, showed decreased Drk level. (C) Ubr1 RNAi did not rescue the decreased Drk level in *psid-D1* clones marked by GFP expression.

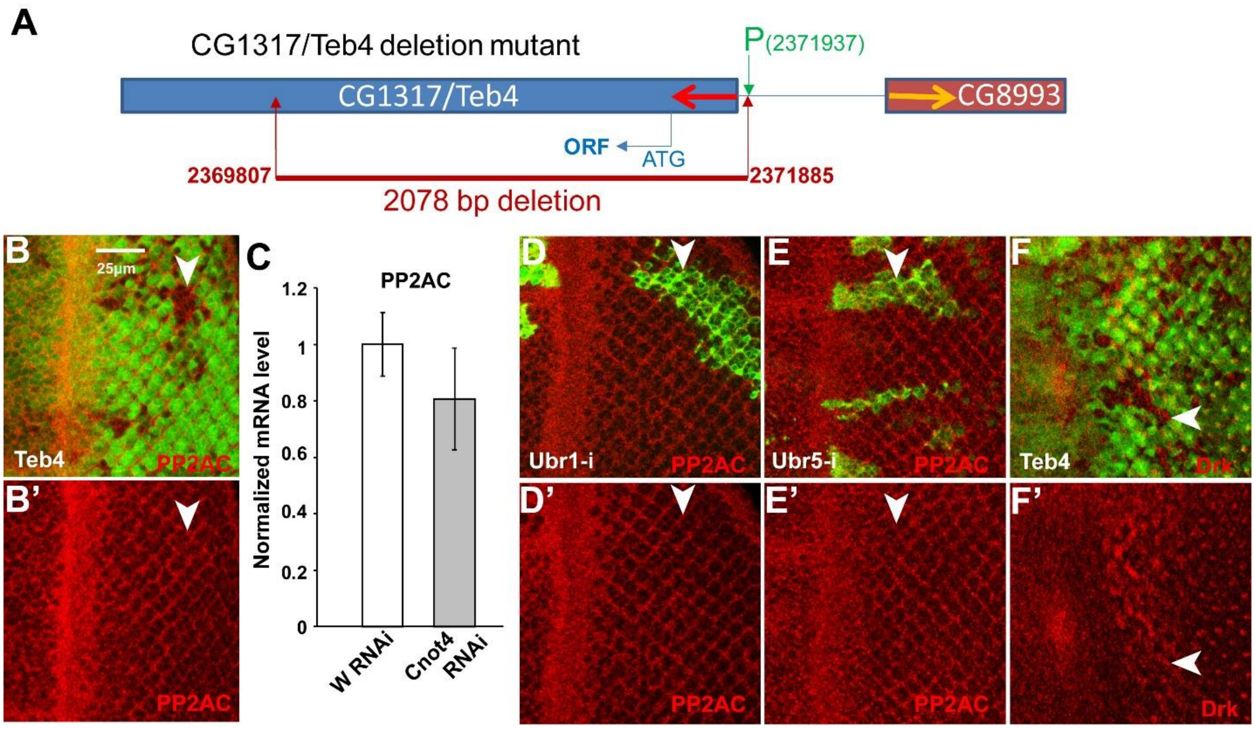


**Fig. S5. PP2AC level was not affected by inactivation of E3 ubiquitin ligases CG1317/Teb4, Ubr1, or Ubr5**

(A) Genomic structure of a deletion allele of *CG1317/Teb4 Δ6-1*, generated from imprecise excision of *CG1317 P* element (BL20646). DNA sequencing data revealed a 2078 bp deletion, which starts from 14 bp upstream of CG1317. The numbers in the diagram indicate the precise location of deletion in the fly genome. (B) *CG1317/Teb4 Δ6-1* mutant clones (pointed by white arrowhead) in eye disc, marked by lack of GFP, did not affect PP2AC levels. (C) Quantitative RT-PCR results from RNA isolated from Cnot4 RNAi or control W RNAi expressing 3$^{rd}$ instar eye/antenna discs. No significant difference was observed in PP2AC mRNA levels. (D-E) Clones of cells (pointed by white arrowheads) with GFP and Ubr1 RNAi (D) or Ubr5 RNAi (E) expression did not affect PP2AC levels in eye discs. (F) *CG1317/Teb4 Δ6-1* mutant clones (pointed by white arrowhead) in eye disc, marked by lack of GFP, did not affect Drk levels.

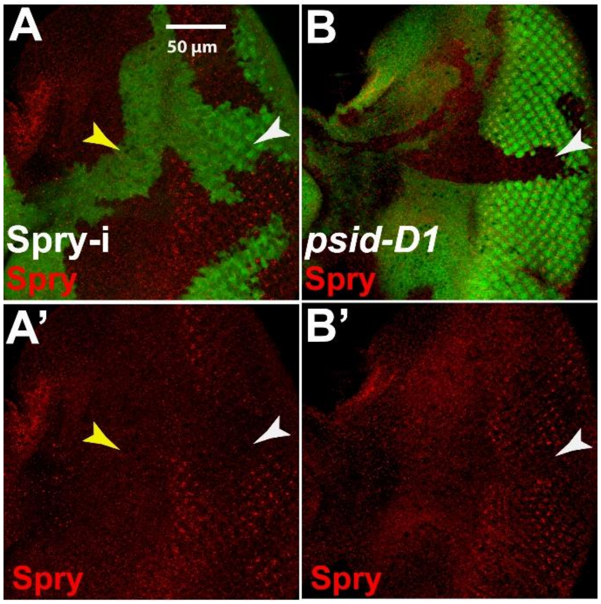


**Fig. S6. Effect of *psid* mutation on Sprouty protein level**. Reduced levels of Spry were observed in clones of cells expressing Spry RNAi (A, RNAi cells were labeled with GFP). White and yellow arrowheads in (A) point to RNAi cells located in the posterior or the anterior region of eye disc. Reduced levels of Spry were also observed in *psid-D1* mutant clones (B, white arrowhead. Mutant clones were marked by absence of GFP). As *psid-D1* mutation reduces EGFR signaling, which is required for Sprouty expression observed in posterior eye discs, these results did not exclude the possibility that *psid* may also affect Spry protein degradation.

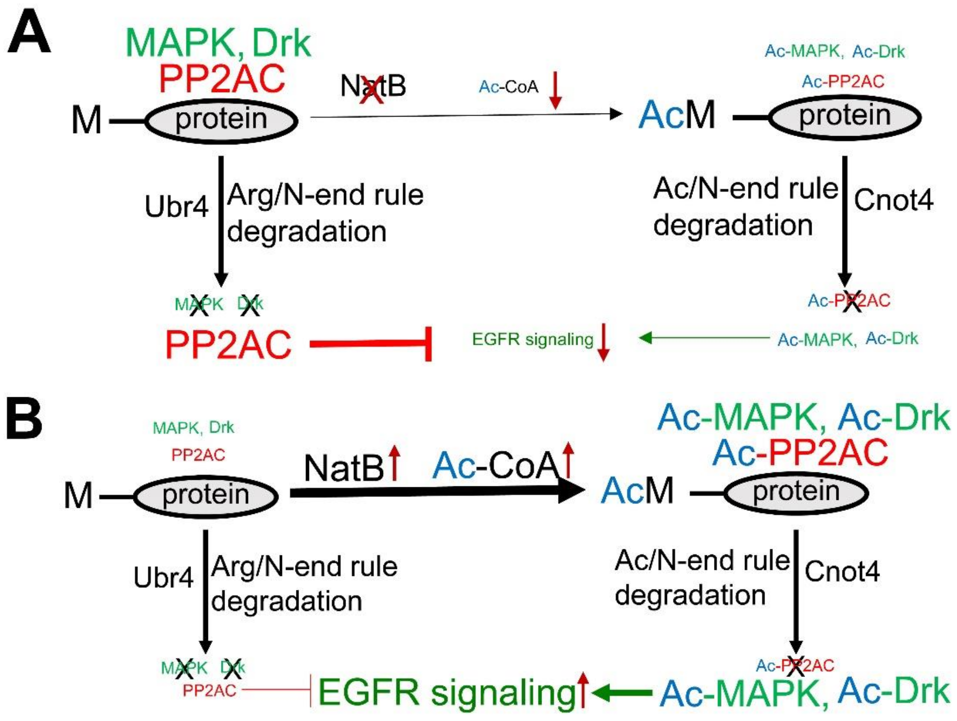


**Fig. S7. Proposed model for the regulation of EGFR signaling by NatB Nt-acetylation and the two branches of N-end rule pathways**. (A) when NatB is inhibited or when Ac-CoA level is low, most Drk, MAPK and PP2AC will not be Nt-acetylated, which results in the inhibition of EGFR signaling due to the accumulation of PP2AC, a negative component of the pathway, and the loss of Drk and MAPK, the positive components of the pathway. (B) When NatB activity is high or when Ac-CoA level is high, more Drk, MAPK, and PP2AC will be Nt-acetylated, which results in the higher levels of EGFR signaling due to the accumulation of acetylated Drk and MAPK and the loss of PP2AC.

**Genotype of flies used**

Fig. S1

w, eyFLP/Y; FRT82B,*Rps3*$^-$, Ubi-GFP/FRT82B, *psid* (*AE*,*EQ, AB, or CJ*)

*rbf*$^{15a\Delta}$,w, eyFLP/Y ; FRT82B, RBF-G3, *Rps3*$^-$, Ubi-GFP/ FRT82B, *psid* (*AE*,*EQ, AB, or CJ*)

eyFLP, Act>CD2>Gal4; UAS-Rbf RNAi / + or UAS-NAA20 RNAi

eyFLP, Act>CD2>Gal4; UAS-NAA20 RNAi

Fig. S2

w, eyFLP /Y; FRT82B, Ubi-GFP / aos-lacz, FRT82B, *psdin* (*AE*,*EQ, psid D1 or D4*)

eyFLP, UAS-Dcr2 / +; CoinFLP-Gal4-UAS-GFP; UAS-NAA20 RNAi

Fig. S3

HsFLP; Act > y >Gal4, UAS-GFP/UAS-Ras$^{v12}$; FRT82B, tub-Gal80/ FRT82B or (FRT82B, *psidD4*)

eyFLP; Act > y >Gal4, UAS-GFP/UAS-Raf$^{gof}$; FRT82B, tub-Gal80/ FRT82B or (FRT82B, *psidD4*)

eyFLP; Act > y >Gal4, UAS-GFP/UAS-EGFR$^{CA}$; FRT82B, tub-Gal80/ FRT82B *scrib* or (FRT82B, *psidD4, scrib*)

eyFLP; Act > y >Gal4, UAS-GFP/ UAS-EGFR$^{CA}$, UAS-psid (WT, SD, or SA); FRT82B, tub-Gal80/ FRT82B, *psidD4, scrib*

Fig. S4

HsFLP; Act > y >Gal4, UAS-GFP; FRT82B, tub-Gal80/ FRT82B, *psid D1 or* (*UAS-Ubr1 RNAi,* FRT82B, *psid D1*)

HsFLP; Act > y >Gal4, UAS-GFP/*Drk-PZ*; FRT82B, tub-Gal80/ FRT82B, *psid D1*

Fig. S5

w, eyFLP; Ubi-GFP, FRT80B/ *Teb4*, FRT80B

HsFLP; tub-Gal80, FRT40A/ FRT40A; Act > y >Gal4, UAS-GFP / UAS-Ubr5 RNAi

HsFLP, Act>CD2>Gal4/Y; UAS-Ubr1 RNAi

Fig. S6

eyFLP, UAS-Dcr2 / +; CoinFLP-Gal4-UAS-GFP;  UAS-Sprouty RNAi

HsFLP; FRT82B,Ubi-GFP / FRT82B, *psid D1*
